# Ingestible capsule for detecting labile inflammatory biomarkers in situ

**DOI:** 10.1101/2022.02.16.480562

**Authors:** ME Inda, M Jimenez, Q Liu, NV Phan, J Ahn, C Steiger, A Wentworth, A Riaz, T Zirtiloglu, K Wong, K Ishida, N Fabian, J Jenkins, J Kuosmanen, W Madani, R McNally, Y Lai, A Hayward, M Mimee, P Nadeau, AP Chandrakasan, G Traverso, RT Yazicigil, TK Lu

## Abstract

Transient molecules in the gastrointestinal (GI) tract, such as nitric oxide and hydrogen sulfide, are key signals and mediators of inflammatory bowel disease (IBD). Because these molecules are extremely short-lived in the body, they are difficult to detect. To track these reactive molecules in the GI tract, we have developed a miniaturized device that integrates genetically-engineered probiotic biosensors with a custom-designed photodetector and readout chip. Leveraging the molecular specificity of living sensors, we genetically encoded bacteria to respond to IBD-associated molecules by luminescing. Low-power electronic readout circuits (nanowatt power) integrated into the device convert the light from just 1 μL of bacterial culture into a wireless signal. We demonstrate biosensor monitoring in the GI tract of small and large animal models and integration of all components into a sub-1.4 cm^3^ ingestible form factor capable of supporting wireless communication. The wireless detection of short-lived, disease-associated molecules could support earlier diagnosis of disease than is currently possible, more accurate tracking of disease progression, and more timely communication between patient and their care team supporting remote personalized care.

## INTRODUCTION

Our ability to diagnose and monitor inflammatory GI disorders would be transformed if we could profile labile, oxidation-related biomarkers and their responses to dietary change and therapies in situ. Many microbiome-related conditions, notably inflammatory bowel disease (IBD), are associated with chronic intestinal inflammation resulting from dysregulated immune homeostasis, specifically, increased oxidation^1^. Malnutrition^2^, antibiotic resistance^3^, antibiotic dysbiosis^4–6^, neurodegenerative diseases^7^, and mitochondrial genetic disorders^8^ are also associated with redox imbalance in the GI tract, and poor responses to chemotherapy^9^ and vaccines^10^, as well as aging^11^, may also be underpinned by oxidative stress.

While the etiology of IBD is not well defined, bacterial infections^12^ and antibiotics^4^ may substantially increase concentrations of oxidants, such as reactive oxygen and reactive nitrogen species (ROS/RNS). These molecules are labile, which can make it difficult to detect their presence or accurately measure their concentration in the body. While there have been reports of devices that sense labile molecules in the GI tract (e.g. oxidizing gases, volatile organic compounds), they are limited to off-the-shelf sensors that use non-specific metal-oxide sensing elements^13,14^. Thus, the current standard of clinical care is limited with respect to our capacity to provide evaluation of the chemical environment underlying the metabolic pathways of both the human host and its resident microbes. Developing non-invasive technologies that can continuously monitor the GI environment in situ would both expand our understanding of what causes inflammation and improve the effectiveness of therapies. Furthermore, diagnosing multifaceted diseases such as IBD, in which biomarker levels vary greatly among patients, would greatly benefit from the simultaneous detection of a panel of oxidation-related biomarkers (e.g. nitric oxide^15^ [NO], ROS^16,17^, thiosulfate^18^ [TS] and tetrathionate^18^ [TT]).

Current methods of diagnosing gastrointestinal (GI) inflammation include (*i*) endoscopy^19^, which is invasive and should only be performed with limited frequency, and (*ii*) stool analysis, which may not accurately reflect intestinal conditions due to differential growth of certain species^20^, ambient oxidation^11^, and loss of labile disease-mediating molecules. Culture enrichment^21^ (e.g. as is done for analyzing low-abundance microorganisms in stool samples with “omics” techniques) may also distort the initial bacterial ratio. Because the fecal microbiota only partially represents the autochthonous microbiota in direct contact with the intestinal mucosa, a biopsy may be required for a complete analysis^22,23^.

Electronic devices that continuously collect, process, and wirelessly transmit information can also be used to analyze the GI tract. However, capsule endoscopy cameras currently approved by the US Food and Drug Administration (FDA)^24^ can not directly measure the molecular mediators of disease, such as ROS/RNS. Other ingestible, ultra-low power electronic devices currently under development can be used to visually evaluate the GI tract and measure gas concentrations, temperature, and pH levels^13,25^ but require functionalization with fragile transducers to convert biochemical information into electronic signals, which limits specificity and robustness^13^.

To overcome these challenges and to leverage the promise of transient biomarker-panels, we combined natural protein-based sensing elements for NO, hydrogen peroxide (H_2_O_2_), TT, and TS with genetically encoded memory circuits, incorporated them into probiotic bacteria, and validated their function in a rodent and porcine model of inflammation. We then integrated these bacterial sensors with a custom-designed integrated photodiode array and readout chip. Our integrated system has a volume below 1.4 cm^3^ and pill form factor conforming with a safe ingestible size for non-deformable dosage forms^26^. This significantly builds on our early prototype^27^ which was only validated for detection of blood in vivo and was considerably larger (>9 cm^3^) than any known safe ingestibles. In addition, our miniaturized wireless bio-electronic pill can safely process data with low-power consumption and transmit it to a portable device such as a smartphone. We have tested this multi-diagnostic pill in pigs, demonstrating that a human-scale diagnostic device can be built to detect transient mediators of GI inflammatory diseases (Fig. 1).

**Fig. 1.**
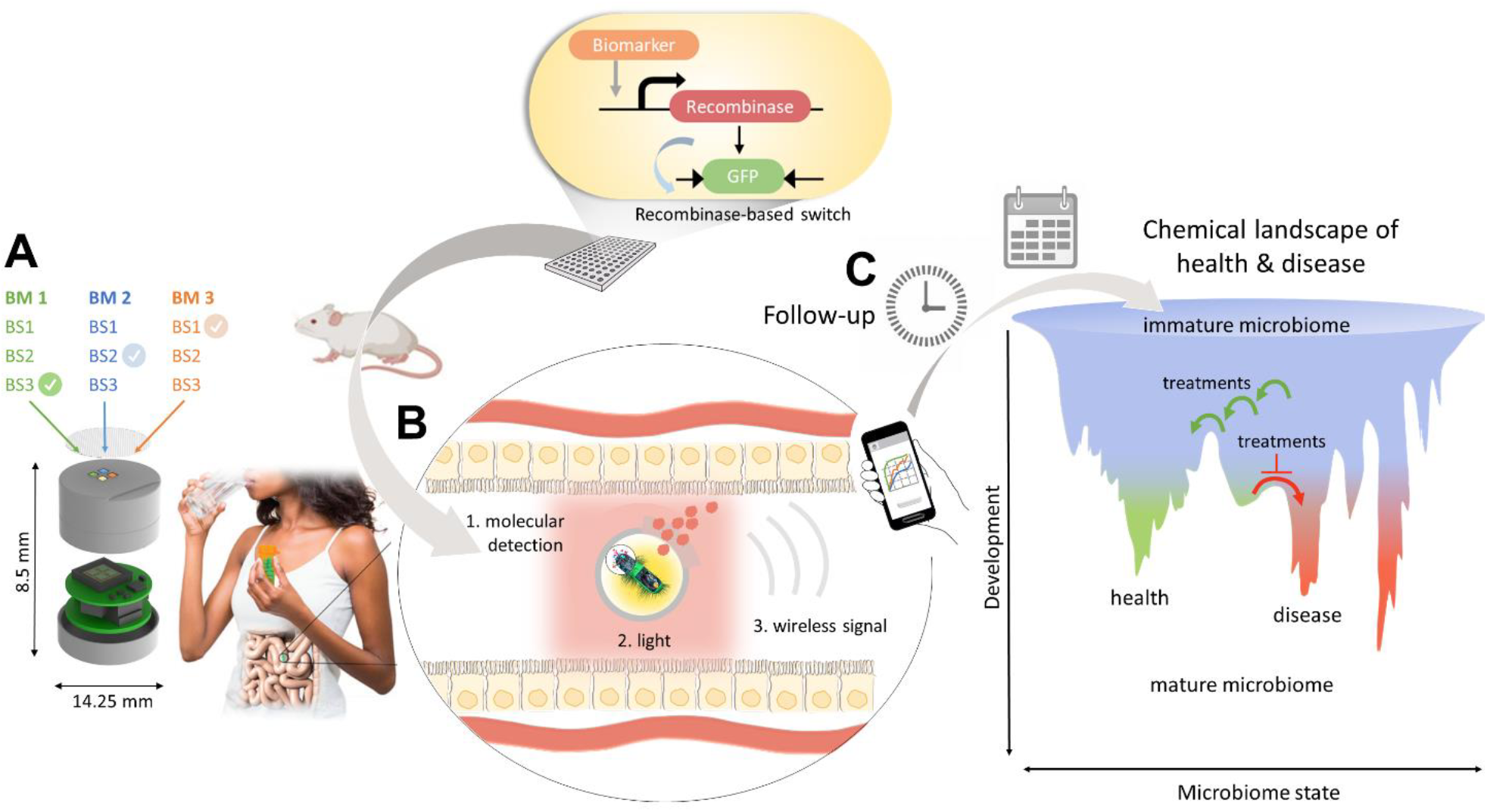
General platform for developing an ingestible capsule for real-time detection of labile mediators of disease. **A.** Probiotic bacteria are engineered to respond to an array of IBD-biomarkers (BM). A recombinase-based genetic memory system is used to validate the bacterial biosensors in animal models. Biosensing bacteria (BS) are then re-engineered to respond by luminescing and packaged in an ingestible capsule along with miniaturized electronics (illustration shows the design and dimension of our fabricated device). **B**. While in transit through the intestines of patients, the biosensing bacteria can sense the metabolites as they are being produced in the body and the integrated ingestible capsule can transmit the bacterial luminescence signal wirelessly to an external device (e.g., a cellular phone). **C**. The device enables remote detection for immediate follow-up after therapy (the clock), as well as monitoring the gut chemical environment for longer-term treatment with personalized therapies or dietary and lifestyle changes (calendar). Microbiome dynamics are depicted at the bottom right, as a simplified model. Starting from infancy, the microbiome gradually reaches one of several adult states, characterized by health or disease. Monitoring the gut chemical environment is essential for timely treatments, since either perturbations that would lead to an unhealthy state can be resisted (red blocked arrow) or the effective treatment plan can be assessed to return to a healthy state (green arrows)^49^. The patient and BS images are reproduced from ref.^50^.

## RESULTS

### Design of biosensing genetic circuits

Intestinal bacteria have natural sensors that continuously detect specific molecules in the gut. Considering the potential immunogenic response to diagnostic microbes, we chose probiotic *Escherichia coli* Nissle 1917 as a chassis due to its resilience in the GI tract and excellent safety profile for long-term use^28,29^ and engineered it to detect the labile IBD-mediating molecules NO, H_2_O_2_, TS, and TT (Fig. S1).

We first created memory circuits that could record NO exposure of bacterial biosensors as they travel through the GI tract and then report any exposure when recovered from feces. Several bacterial NO sensors control the expression of NO reductases, which detoxify NO inside the gut^30^. We chose the sensor NorR because it differentiates NO from other reactive nitrogen species (NOx)^20,31,32^ that are abundant in the gut environment. NorR activates transcription from the norV promoter (PnorV).

Recombinases recognize specific DNA sequences and can invert them, leaving long-lasting changes in DNA^33^. To create memory circuits to report NO exposure, we combined a bacterial NO biosensor with a DNA recombinase core circuit (Fig. 2A). Once an exposure is recorded, the information is stored in the DNA of the bacteria and passed from generation to generation. The information can then be retrieved by measuring green fluorescent protein (GFP) expression.

**Fig. 2.**
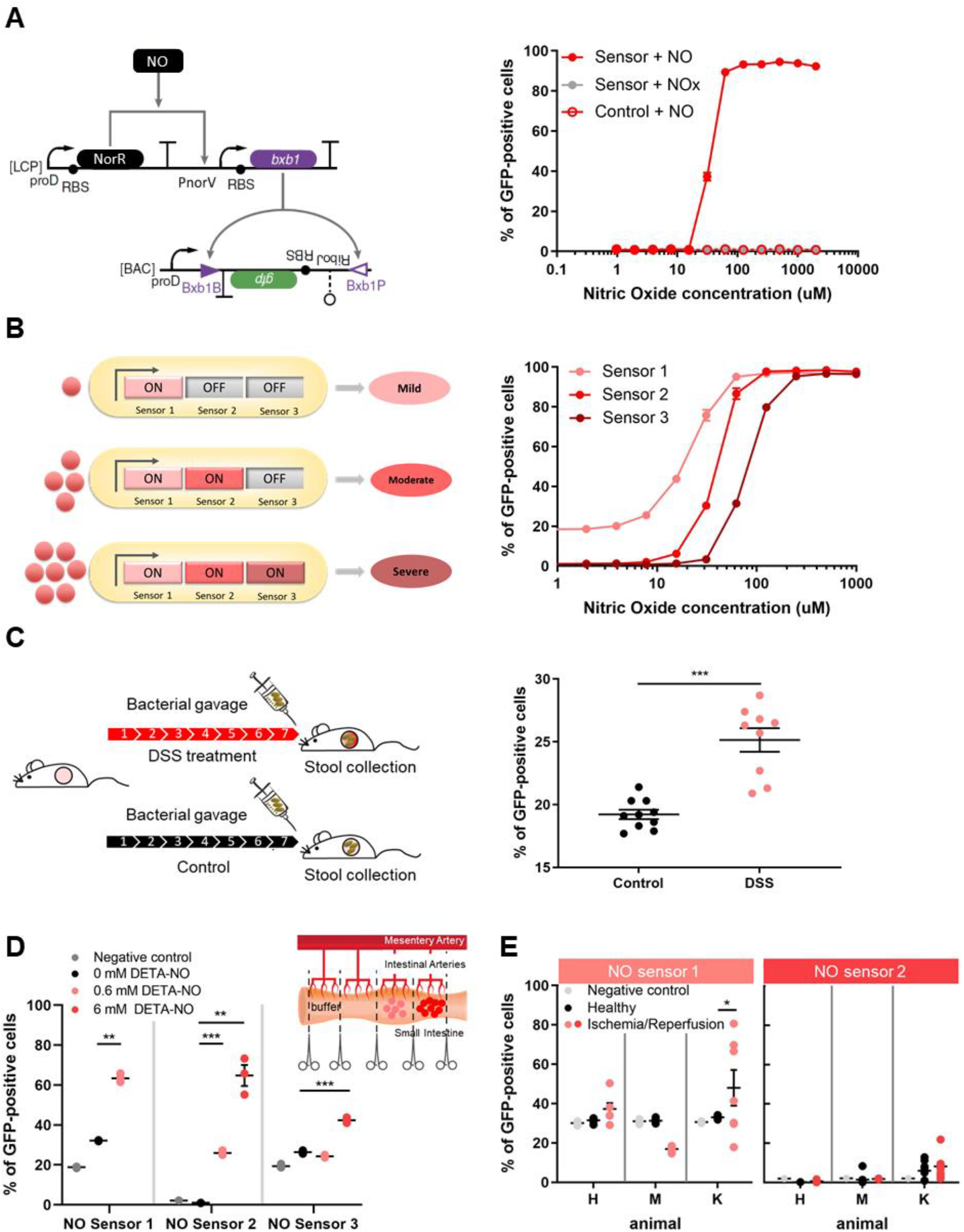
Validation in vitro and in vivo of probiotic bacteria engineered to detect nitric oxide (NO). **A.** To sense NO, the transcription factor NorR, which activates the PnorV promoter, was constitutively expressed from a low copy number plasmid (LCP, left panel). In response to NO, NorR binds to the PnorV promoter, activating the transcription of bxb1 recombinase, leading to inversion of the GFP expression cassette (located between attB and attP sites, triangles, in a bacterial artificial chromosome, BAC) and expression of GFP. Flow cytometry was used to determine the percent of GFP-positive cells at various NO concentrations (right panel). Other reactive nitrogen species (NOx) did not activate the system. **B.** The NO biosensor was evaluated for its use as an NO disease stage detector: NorR, constitutively expressed from a library of ribosome-binding sites (RBS), exhibited different NO activation thresholds. Selected NO sensors detected three concentrations (15, 30, and 80uM), which could correspond respectively to mild, moderate, and severe states of inflammation^51^. Based on the recombinase system described in (A), flow cytometry was used to measure the percentage of GFP-positive cells at different concentrations of NO (right panel). For each point, the mean of three biological replicates, each with n=10,000 flow cytometry events is plotted, and the error bars are the standard error of the mean (SEM). **C.** The NO biosensor was validated in murine models of chemically-induced colitis. Experimental timeline (left panel): C57BL/6 male mice were treated for 7 days with 3% DSS in drinking water and gavaged every other day with engineered *E. coli* carrying the NO memory system. Stool was collected 6 hours post-gavage. Control mice (no DSS treatment) were simultaneously gavaged with the same engineered bacterial strain. The sensor for NO was significantly activated on day 6 of DSS treatment (right panel). Eight-week-old C57BL/6 male mice were used for this study. ‘‘DSS” samples, n = 10, “Control” samples, n =10. ***p < 0.001, Student’s t-test. **D.** The NO sensor was validated in pigs. Experimental design (inset, left panel): intestines were clamped to separate the different compartments to test control vs treated compartment. Bacterial sensors were placed in the different compartments (left panel). NO Sensor 1 was significantly activated in the presence of 0.6mM DETA-NO. NO Sensor 3 only detected 6 mM DETA-NO. NO Sensor 2, could detect the intermediate range, from 0.6 to 6mM. **E.** Ischemia/Reperfusion model of inflammation in pigs. For (D-E) the bacteria were collected from the intestine after two hours of exposure to the analyte or the ischemia/reperfusion, and the percent of GFP-positive cells was measured by flow cytometry (n=10,000 events) from separately grown culture post retrieval. Lines represent the mean of these replicates and error bars are the SEM. Here, we show data of three independent experiments (three animals [H, M, K] on different days, multiple compartments per animal). *p<0.5, **p<0.01, ***p<0.001, Student’s t test.

Using flow cytometry, we measured GFP expression and compared it to the concentration of DETA/NO (diethylenetriamine/nitric oxide adduct). The percentage of cells that were GFP ‘ON’, (% GFP - positive cells)^16^ was calculated at each concentration of NO. After multiple rounds of optimization to improve the signal-to-noise ratio (SNR) of the genetic circuits and controls (Fig. S2A-C), our NO recombinase-based memory system responded to a concentration threshold of 30 μM diethylenetriamine/nitric oxide (DETA/NO) and was not influenced by NOx compounds present in the diet, showing high specificity for NO (Fig. 2A). The performance of the NO sensor was tested under anaerobic conditions (Fig. S2D) and over time (Fig. S2E). Similarly, we built recombinase-based memory circuits for ROS, TS, and TT detection (Fig. S3A).

We also created a disease stage detector (DSD) based on the concentration of NO detected, in which each stage could indicate a different level of inflammation - mild, moderate and severe (Fig. 2B). The incorporated recombinase-based switch was also used to discretize the biomarker input levels and create digital memories in the cells^16^. Tuning the sensitivity of the NO biosensor resulted in different activation thresholds, so that physiologically relevant concentrations of the biomarker (low, medium, or high) could be measured in animal models. All of the sensors developed for in vivo validation in mice measured GFP expression as a readout of the memory system activation; the sensors were activated within minutes (Fig. S2E).

### Disease detection in animal models of IBD via bacterial sensors

To examine functionality in vivo, we first evaluated whether bacterial sensors passing through the GI tract could detect chemically-induced GI inflammation in a mouse model of colitis (Fig. 2C). After inflammation was induced with dextran sodium sulfate (DSS), control and treated mice were orally gavaged with the NO biosensors. After six hours^27^, the percentage of GFP+ cells recovered from fecal samples was analyzed by flow cytometry. NO biosensors demonstrated significantly more GFP+ activation in fecal pellets from treated mice than in those from healthy controls (Fig. 2C). Our novel biosensor design thus was able to detect the presence of NO as a marker of GI inflammation in vivo. Inflammation in the colitis model was measured over time (Fig. S4A) and independently validated (Fig. S4B-C).

GFP activation increased significantly at day nine after the start of DSS treatment (Fig. S4A) correlating with the peak of nitric oxide synthase (iNOS) activation^34^. Tracking NO with the DSD revealed an exacerbated inflammatory response following antibiotic treatment in a chronic inflammation model (Fig. S4D). We also tested our bacterial sensors for ROS (H_2_O_2_), TT and TS; activation was detected as inflammation progressed, with TS detection at day six after the start of DSS treatment and TT (TS product of oxidation) at day eight, which overlapped with ROS detection (Fig. S3B-C).

To validate the bacterial sensors in a disease model comparable in physiology and scale to human anatomy^35^, we also tested the bacterial biosensors in pigs via an ischemia/reperfusion injury model of intestinal inflammation^36^. We injected the bacterial biosensors directly into the intestinal lumen of a sedated animal, in either inflamed segments, or healthy segments with or without added biomarkers. The NO sensing bacteria could detect different concentrations of NO in the control-treated group (Fig. 2D) and lead to a positive signal in the disease group (ischemia/reperfusion, Fig. 2E); the ROS, TS, and TT sensors also detected significant quantities of analytes in control-treated pigs (Fig. S4E).

### Validation of an integrated, sub-1.4 mL, ingestible bacterial-electronic pill

To advance the diagnostic potential of the bacterial sensors towards clinical application, we next integrated them into a bacterial-electronic pill. Specifically, we sought to design a system with a size and form factor conforming to a proven non-deformable dosage form (Fig. S5). To achieve this goal, we developed an optimized luminescent readout, a custom microelectronic bioluminescence detector and bacterial chambers built directly into the pill casing.

Our multi-diagnostic device requires nutrients and analytes to be exchanged efficiently while retaining the engineered microbes and simultaneously allowing the generated light signal to reach the electronic detectors. To meet this design requirement, we developed a pill casing that incorporates a bacterial-electronic chamber interface in a unibody design (Fig. 3A-B). This small tablet-shaped housing precisely aligns the bacteria in a two by two array with the microelectronic photodiodes while maintaining a hermetic seal around the electronic components. This design enables sensing from an array of engineered microbial strains designed to express luciferase in response to several biomarkers (i.e., NO, ROS, TT and negative reference). The integrated chambers (Fig. S6) are sealed with porous membranes (nominal pore size, 0.4 μm) to retain the bacterial cells. The porous membranes, when placed in feces, did not interfere with the detection of the target molecules (Fig. S7).

**Fig. 3.**
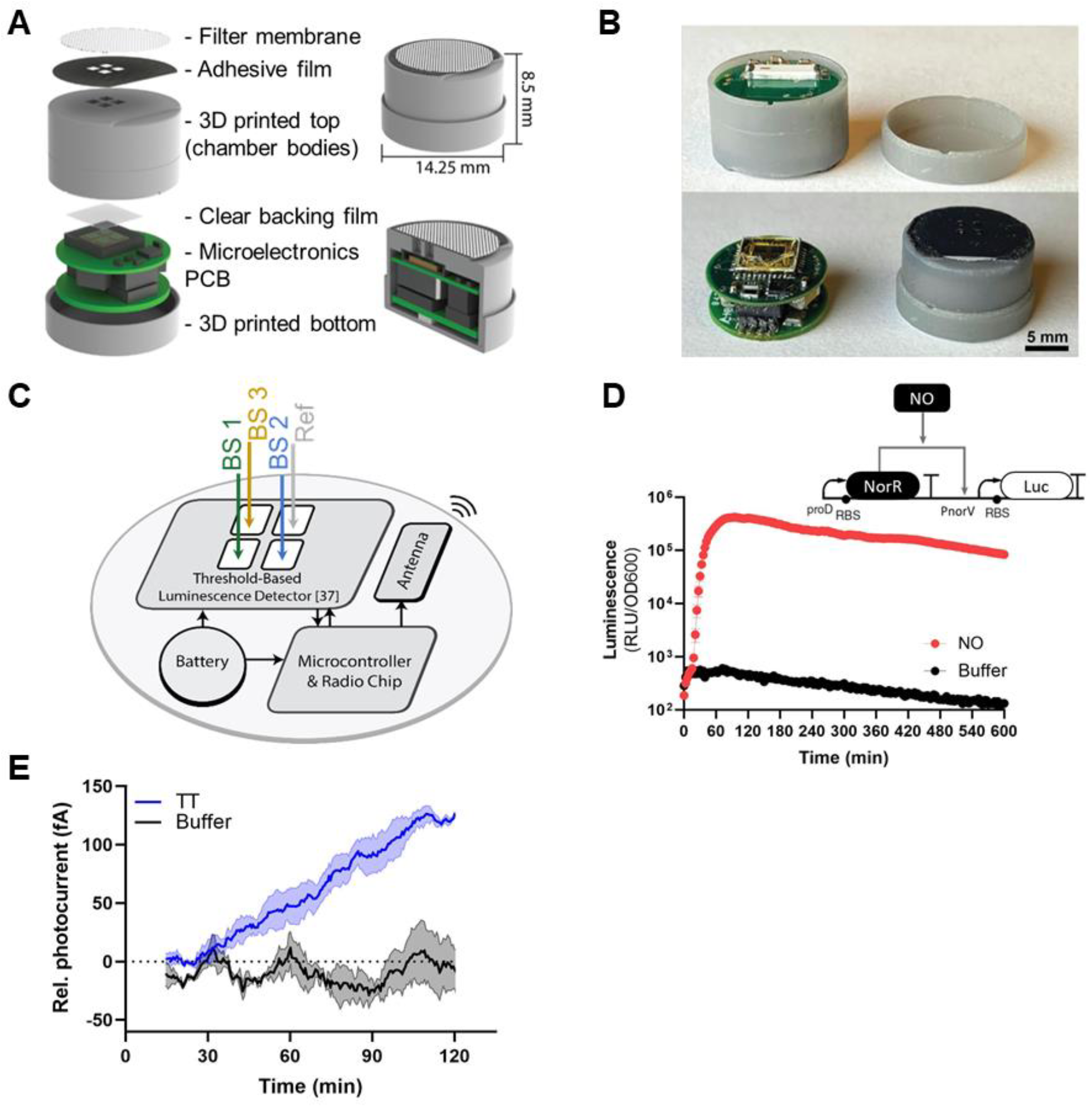
Design and in vitro characterization of the device for miniaturized wireless sensing with cell-based biosensors. **A**. Basic components and dimensions of the device. **B.** Design of a miniaturized pill casing with a bacterial-electronic chamber interface. Photos showing: (top) side view of the device; (bottom) fully assembled pill with the permeable membrane attached. The bacterial chamber/casing unibody design uses a thin clear backing film to place the bacteria as close to the photosensitive electronic chip as possible. A double-sided adhesive film enables a low-profile seal to the permeable outer filter membrane. Scale bar = 5 mm. **C.** The outputs of multiple biosensors (BS) intended to be studied together can be measured with this array (for example, the H_2_O_2_, TS, and TT sensors).The detailed schematic of microelectronics PCB is shown in Fig. S10. A threshold-based bioluminescence detector with a CMOS-integrated photodiode^37^ array was used to detect bacterial sensor output. **D.** Genetic circuit and kinetic response of NO sensors in bacterial growth media supplemented with 3 mM NO; RLU, relative luminescence units. Error bars represent SEM of three independent biological replicates. **E.** Wireless signal over time from the TT sensor encapsulated in the device and immersed in bacterial growth media supplemented with 100 mM TT. Low-power CMOS-integrated photodiodes converted bioluminescence emitted from the bacterial sensor into a photocurrent, which was converted into quantifiable digital data and transmitted wirelessly to the external device. Lines represent the mean, and error bars denote the SEM for three independent replicates conducted with one induced device (TT) and one uninduced device (buffer); Rel. photocurrent, Relative Photocurrent.

To generate bacterial sensors compatible with the photosensitive electronics, we replaced the GFP readout with a self-contained bioluminescence readout (i.e. the luxCDABE operon from *Photorhabdus luminescens*)^27^. We constructed this genetic circuit in *E. coli* Nissle 1917 and exposed the resultant strain to DETA/NO as a source of NO. The NO biosensing bacteria responded rapidly to DETA/NO exposure (t_max_ = 60 mins) with a high luminescence output and a SNR of 170 (Fig. 3C). Luminescence production was also induced by DETA/NO under anaerobic conditions (Fig. S8). The other inflammatory sensors were also built and characterized in vitro (Fig. S9A-C), with TT sensor reaching the highest luminescence values in simulated intestinal fluid (Fig. S9B).

Lastly, we designed a millimeter-scale capsule printed circuit board (PCB) housing a complementary metal-oxide-semiconductor (CMOS) bioluminescence detector chip with an integrated photodiode array achieving high sensitivity (Fig. 3A). The custom chip integrates a threshold-based bioluminescence detector, time-to-digital converter, voltage references, voltage regulators, and four 1 mm by 1 mm CMOS-compatible photodiodes^24^ (Fig. S10)^37^. In addition to the custom chip, the ingestible capsule also includes a commercial microcontroller and wireless transmitter. Bioluminescence from activated cells was detected by CMOS-integrated photodiodes located below each chamber. Custom-designed electronics processed the luminescence data by periodically sampling the photon-generated charge (with a programmable integration time of ~26s). The detected luminescence was converted to a digital code by the low-power luminescence readout chip and transmitted wirelessly for calibration, display, and recording.

When tested in vitro, using 1 μL of sensor bacteria culture per chamber, the integrated device successfully detected the presence of TT. We recorded a net 175 fA photocurrent produced by the induced TT bacteria sensor, with a baseline of ~0 fA from the uninduced control (Fig. 3D).

The integrated device was also tested in vivo in pig small intestines as a model for the complex milieu of the GI tract. Upon induction, luminescence was detected by the custom-designed electronic readout circuits in the capsule; the information was wirelessly transmitted in real time from inside the body of the pig to an external recording device (Fig. 4A-B). This design allowed remote monitoring of biomolecules in the gut for four hours, with 75 fA relative photocurrent detected in the induced compartment, and a baseline of −90 fA from the uninduced control (Fig. 4C and Fig. S11). The receiver operating characteristic of the TT sensing reaches a sensitivity and specificity of 100% at 60 min. The miniaturized device can thus detect small amounts of analyte in the harsh intestinal environmen with high specificity and sensitivity (Fig. 4D and Fig. S12).

**Fig. 4.**
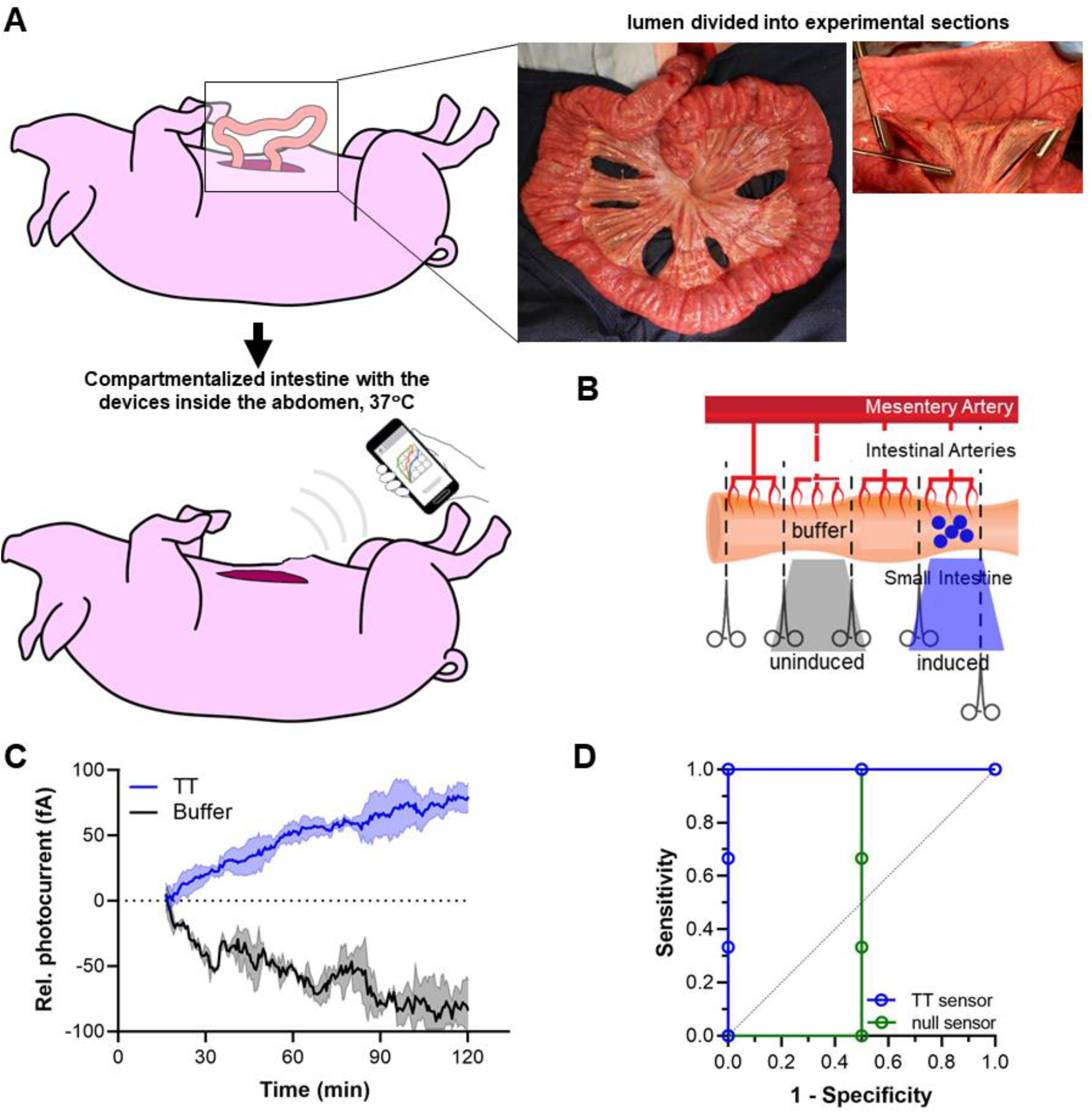
Validation of the whole integrated device for miniaturized wireless biosensing in pigs. **A.** Schematic of the experimental flow, the bacterial biosensors encapsulated in the device were tested in situ in a terminal procedure in swine, for which anesthetized Yorkshire pigs of around 90 kg were used. After opening the abdominal cavity, we placed the device through a small incision directly into the lumen of the pig’s small intestine, compartmentalized with clamps. Compartmentalized intestines were kept inside the abdomen, at 37°C. Wireless signals transmitted from inside the abdomen were detected by a commercial receiver, which is connected to a device, such as a laptop computer or, alternatively, by a cellular phone. **B.** Schematic diagram of compartmentalization of a section of the intestine clamped for experimentation. (1) TT (100 mM) was injected with a syringe into the clamped intestinal compartment. Buffer was added as a control, in another compartment. **C.** Kinetics of TT sensor embedded in the abdominal cavity of the pig. The response of the device placed in the compartment with TT was clearly distinguishable from that of the device in the compartment with the buffer control. Dark lines represent the mean of independent experiments in different pigs with up to two devices in separate compartments (+TT and +buffer). Shading represents the SEM. ‘‘TT” measurements, n = 3 and “Buffer” measurements, n =2. *P < 0.05, Student’s t test; Rel. photocurrent, Relative Photocurrent. **D.** Receiver operating characteristic (ROC) of the device sensing over time. Perfect detection is achieved at t = 60 min.

## DISCUSSION

We have built an ingestible microbial biosensor that can sense an array of biomarkers in situ, as they are being produced. This technology can be applied as a tool to support remote disease management by providing quantitative, real-time, and multiplexed information linking GI tract microbiome perturbations to disease. To validate the cell-based biosensors in preclinical disease models, we coupled engineered sensing bacteria with a recombinase-based memory system. Our memory system records information as soon as metabolites are produced in the gut, activating switches within minutes of exposure. The recombinase-based switch discretizes the magnitude of a given biomarker; this quantitative response may align with disease stage and therefore indicate severity of inflammation. Although some of the parameters that can be measured with this device may have no absolute “healthy range”, measurements taken over time would reveal patterns predictive of acute disease episodes (flares); disease symptoms could then be anticipated. Similarly, the switch acts as a peak detector for sensing and recording maximum levels of intestinal biomarkers.

To create our miniaturized low-power ingestible electronic device, we integrated CMOS-compatible photodiodes with discrete-time signal processing circuits, which made it possible to miniaturize the whole capsule to a size below 1.4 cm^3^ and simultaneously detect multiple disease biomarkers on a stringent power budget. This integrated CMOS system combined with our unibody chamber pill casing design allowed us to detect the bioluminescent signal from just 1 μL of bacterial culture in the milieu of the intestinal lumen. In addition, the coin-cell battery can power the ingestible capsule for a month^37^, so this device could also potentially be used as an implant^38^.

The diagnostic accuracy and specificity of our device are based on the simultaneous testing of an array of labile by-products of inflammation (e.g., NO and ROS), intestinal gases (e.g., H_2_S measured as TS), and microbiome-derived biomolecules (e.g., TT). As biomarker levels may vary greatly among patients, a panel of biomarkers would be required to accurately diagnose IBD and other multi-faceted diseases.

Our ingestible device offers a route for non-invasively evaluating changes in the intestinal biochemical milieu and overcomes the limitations of microbiome characterization by 16S rRNA or metagenomic sequencing^39,40^, as well as current research applications of ingestible biosensors in animal models, which require the complex analysis of bacterial gene expression or RNA/DNA in stool^18,41–45^. The biosensors described here also have the potential to expand on the range of biomarkers being targeted by other ingestible electronic systems^46^. The capsule could be designed to report its location while in transit^47^ and perform cell-based computation to further expand the multiplex capabilities of the underlying electronic system. For example, AND-gates^48^ could be incorporated to determine the co-localization of biomarkers for understanding metabolic pathways or biomarker discovery in animal models for many microbiome-linked diseases.

This device can be developed as a first-line at-home screen for non-invasive continuous monitoring of the chemical environment of the GI tract and customized for numerous GI disorders, thus offering a safe and inexpensive point-of-care alternative to imaging capsules for endoscopy. Tracking and quantitatively assessing multiple biomarkers can potentially provide a framework for patients to assess the effects of diet, lifestyle, and other interventions that require routine screening to improve health outcomes.

## Author contributions

MEI, MJ, QL, NP, CS, MM, PN, GT, RTY, TLK conceived and designed the research; MEI, MJ, QL, CS, AW, MM, PN, AC, GT, RTY, TLK conceptualized the miniaturized pill form-factor, including integration of the bacteria, electronics and pill casings; MEI designed and performed in vitro biological experiments; QL, AR, TZ designed and built the integrated electronic circuits; MEI designed and performed in vivo mouse experiments; MJ, JA, AW, KW developed the pill casing manufacturing process and validated the pill casing robustness in vitro, including membrane attachment; MJ, AW, KW, RM generated 3D-printed device components. MEI, MJ, QL, NP, CS, AH, GT validated early prototypes; MJ, NP, CS, KI, JJ, JK, AH, GT conceptualized and validated the intestinal compartment animal model; KI, FN, JJ, JK, AH carried out animal husbandry and anesthesia of swine. MEI, MJ, QL experimentally tested the function of the integrated devices in vitro and in vivo; MEI, MJ, QL performed formal analysis of the data; MEI, MJ, QL, JA, PN, RTY contributed to data analysis of in vitro and in vivo experiments of the integrated devices; MEI, MJ, QL, NP, JA, AW, PN, RTY contributed to visualization; MEI, MJ, QL, MM, PN, AC, GT, RTY, TKL wrote the manuscript; MEI, MJ, QL, PN, CS, AW, AH, GT, RTY managed daily project progress and personnel. MEI, TKL supervised and managed general project administration; RTY, GT supervised and managed funding of the project related to the integrated electronics and integrated pill casings and swine husbandry, respectively; MEI, MJ, NP, CS, YL, MM, PN, AC, RTY, GT, TLK, contributed with funding acquisition.

## Funding

Supported by Leona M. and Harry B. Helmsley Charitable Trust (3239), Pew Charitable Trusts (to M.E. Inda; 00030623). MJ was supported by the Translational Research Institute of Space Health through Cooperative Agreement NNX16AO69A. GT was supported in part by the Department of Mechanical Engineering, MIT and the Karl van Tassel (1925) Career Development Professorship, MIT.

The patient and BS images in Fig 1 and FS1 are reproduced from *The Journal of allergy and clinical immunology* **144**, 645–647. Inda, M. E., Mimee, M. & Lu, T. K. “Cell-based biosensors for immunology, inflammation, and allergy”. Copyright (2019), with permission from Elsevier.

## References

1. Bourgonje, A. R. et al. Oxidative Stress and Redox-Modulating Therapeutics in Inflammatory Bowel Disease. Trends Mol. Med. 26, 1034–1046 (2020).

2. Million, M. et al. Increased Gut Redox and Depletion of Anaerobic and Methanogenic Prokaryotes in Severe Acute Malnutrition. Sci. Rep. 6, 26051 (2016).

3. Pribis, J. P. et al. Gamblers: An Antibiotic-Induced Evolvable Cell Subpopulation Differentiated by Reactive-Oxygen-Induced General Stress Response. Mol. Cell 74, 785–800.e7 (2019).

4. Reese, A. T. et al. Antibiotic-induced changes in the microbiota disrupt redox dynamics in the gut. Elife 7, 1–22 (2018).

5. Rivera-Chávez, F., Lopez, C. A. & Bäumler, A. J. Oxygen as a driver of gut dysbiosis. Free Radic. Biol. Med. 105, 93–101 (2017).

6. Rivera-Chávez, F. et al. Depletion of Butyrate-Producing *Clostridia* from the Gut Microbiota Drives an Aerobic Luminal Expansion of *Salmonella*. Cell Host Microbe 19, 443–454 (2016).

7. Dumitrescu, L. et al. Oxidative Stress and the Microbiota-Gut-Brain Axis. Oxid. Med. Cell. Longev. 2018, 2406594 (2018).

8. Yardeni, T. et al. Host mitochondria influence gut microbiome diversity: A role for ROS. Sci. Signal. 12, eaaw3159 (2019).

9. Jose, S., Bhalla, P. & Suraishkumar, G. K. Oxidative stress decreases the redox ratio and folate content in the gut microbe, Enterococcus durans (MTCC 3031). Sci. Rep. 8, 12138 (2018).

10. Hagan, T. et al. Antibiotics-Driven Gut Microbiome Perturbation Alters Immunity to Vaccines in Humans. Cell 178, 1313–1328.e13 (2019).

11. Million, M. & Raoult, D. Linking gut redox to human microbiome. Hum. Microbiome J. 10, 27–32 (2018).

12. Rivera-Chávez, F. & Bäumler, A. J. The Pyromaniac Inside You: Salmonella Metabolism in the Host Gut. Annu. Rev. Microbiol. 69, 31–48 (2015).

13. Kalantar-Zadeh, K. et al. A human pilot trial of ingestible electronic capsules capable of sensing different gases in the gut. Nat. Electron. 1, 79–87 (2018).

14. Steiger, C. et al. Dynamic Monitoring of Systemic Biomarkers with Gastric Sensors. Adv. Sci. (Weinheim, Baden-Wurttemberg, Ger. 8, e2102861 (2021).

15. Archer, E. J., Robinson, A. B. & Süel, G. M. Engineered E. coli that detect and respond to gut inflammation through nitric oxide sensing. ACS Synth. Biol. 1, 451–457 (2012).

16. Rubens, J. R., Selvaggio, G. & Lu, T. K. Synthetic mixed-signal computation in living cells. Nat. Commun. 7, 1–10 (2016).

17. Müller, I. E. et al. Gene networks that compensate for crosstalk with crosstalk. Nat. Commun. 10, 1–8 (2019).

18. Daeffler, K. N. et al. Engineering bacterial thiosulfate and tetrathionate sensors for detecting gut inflammation. Mol. Syst. Biol. 13, 923 (2017).

19. Annese, V. et al. European Evidence-based Consensus: Inflammatory Bowel Disease and Malignancies. J. Crohns. Colitis 9, 945–965 (2015).

20. Amir, A. et al. Room-temperature, Correcting for microbial blooms in fecal samples during shipping. mSystems 2, e00199–16 (2017).

21. Raymond, F. et al. Culture-enriched human gut microbiomes reveal core and accessory resistance genes. Microbiome 7, 56 (2019).

22. Mangifesta, M. et al. Mucosal microbiota of intestinal polyps reveals putative biomarkers of colorectal cancer. Sci. Rep. 8, 1–9 (2018).

23. Jain, U. et al. Debaryomyces is enriched in Crohn’s disease intestinal tissue and impairs healing in mice. Science (80-.). 371, 1154–1159 (2021).

24. Swain, P. Wireless capsule endoscopy. 48–50 (2003).

25. van der Schaar, P. J. et al. A novel ingestible electronic drug delivery and monitoring device. Gastrointest. Endosc. 78, 520–528 (2013).

26. Bass, D. M., Prevo, M. & Waxman, D. S. Gastrointestinal Safety of an Extended-Release, Nondeformable, Oral Dosage Form (OROS®)1. Drug Saf. 25, 1021–1033 (2002).

27. Mimee, M. et al. An ingestible bacterial-electronic system to monitor gastrointestinal health. Science (80-.). 360, 915–918 (2018).

28. Isabella, V. M. et al. Development of a synthetic live bacterial therapeutic for the human metabolic disease phenylketonuria. Nat. Biotechnol. 36, 857–864 (2018).

29. Kurtz, C. et al. Translational Development of Microbiome-Based Therapeutics: Kinetics of E. coli Nissle and Engineered Strains in Humans and Nonhuman Primates. Clin. Transl. Sci. 11, 200–207 (2018).

30. Rodionov, D. A., Dubchak, I. L., Arkin, A. P., Alm, E. J. & Gelfand, M. S. Dissimilatory metabolism of nitrogen oxides in bacteria: comparative reconstruction of transcriptional networks. PLoS Comput. Biol. 1, e55 (2005).

31. Bush, M., Ghosh, T., Tucker, N., Zhang, X. & Dixon, R. Transcriptional regulation by the dedicated nitric oxide sensor, NorR: A route towards NO detoxification. Biochem. Soc. Trans. 39, 289–293 (2011).

32. Tucker, N. P. et al. Analysis of the nitric oxide-sensing non-heme iron center in the NorR regulatory protein. J. Biol. Chem. 283, 908–918 (2008).

33. Ceze, L., Nivala, J. & Strauss, K. Molecular digital data storage using DNA. Nat. Rev. Genet. 20, 456–466 (2019).

34. Beck, P. L. et al. Paradoxical roles of different nitric oxide synthase isoforms in colonic injury. Am. J. Physiol. - Gastrointest. Liver Physiol. 286, 137–147 (2004).

35. Jiminez, J. A., Uwiera, T. C., Douglas Inglis, G. & Uwiera, R. R. E. Animal models to study acute and chronic intestinal inflammation in mammals. Gut Pathog. 7, 29 (2015).

36. Strand-Amundsen, R. J. et al. Ischemia/reperfusion injury in porcine intestine - Viability assessment. World J. Gastroenterol. 24, 2009–2023 (2018).

37. Liu, Q. et al. Zero-Crossing-Based Bio-Engineered Sensor. in 2021 IEEE Custom Integrated Circuits Conference (CICC) 1–2 (2021). doi:10.1109/CICC51472.2021.9431409

38. Liu, X. et al. Magnetic Living Hydrogels for Intestinal Localization, Retention, and Diagnosis. Adv. Funct. Mater. 31, 2010918 (2021).

39. Vujkovic-cvijin, I. et al. Host variables confound gut microbiota studies of human disease. Nature 587, (2020).

40. Chiu, C. Y. & Miller, S. A. Clinical metagenomics. Nat. Rev. Genet. 20, 341–355 (2019).

41. Kotula, J. W. et al. Programmable bacteria detect and record an environmental signal in the mammalian gut. Proc. Natl. Acad. Sci. 111, 4838–4843 (2014).

42. Riglar, D. T. et al. Engineered bacteria can function in the mammalian gut long-term as live diagnostics of inflammation. Nat. Biotechnol. 35, 653–658 (2017).

43. Lim, B., Zimmermann, M., Barry, N. A. & Goodman, A. L. Engineered Regulatory Systems Modulate Gene Expression of Human Commensals in the Gut. Cell 169, 547–558.e15 (2017).

44. Mimee, M., Tucker, A. C., Voigt, C. A. & Lu, T. K. Programming a Human Commensal Bacterium, Bacteroides thetaiotaomicron, to Sense and Respond to Stimuli in the Murine Gut Microbiota. Cell Syst. 1, 62–71 (2015).

45. Pickard, J. M. et al. Rapid fucosylation of intestinal epithelium sustains host–commensal symbiosis in sickness. Nature 514, 638–641 (2014).

46. Steiger, C. et al. Ingestible electronics for diagnostics and therapy. Nat. Rev. Mater. 4, 83–98 (2019).

47. Ma, Y., Selby, N. & Adib, F. Minding the billions: Ultra-wideband localization for deployed RFID tags. Proc. Annu. Int. Conf. Mob. Comput. Networking, MOBICOM Part F1312, 248–260 (2017).

48. Siuti, P., Yazbek, J. & Lu, T. K. Synthetic circuits integrating logic and memory in living cells. Nat. Biotechnol. 31, 448–452 (2013).

49. Lloyd-Price, J., Abu-Ali, G. & Huttenhower, C. The healthy human microbiome. Genome Med. 8, 51 (2016).

50. Inda, M. E., Mimee, M. & Lu, T. K. Cell-based biosensors for immunology, inflammation, and allergy. The Journal of allergy and clinical immunology 144, 645–647 (2019).

51. Reinders, C. I. et al. Rectal mucosal nitric oxide in differentiation of inflammatory bowel disease and irritable bowel syndrome. Clin. Gastroenterol. Hepatol. 3, 777–783 (2005).

52. Gibson, D. G. et al. Enzymatic assembly of DNA molecules up to several hundred kilobases. Nat. Methods 6, 343–345 (2009).

