## Supplementary Materials for "Ingestible capsule for detecting labile inflammatory biomarkers in situ"

### SUPPLEMENTARY FIGURES

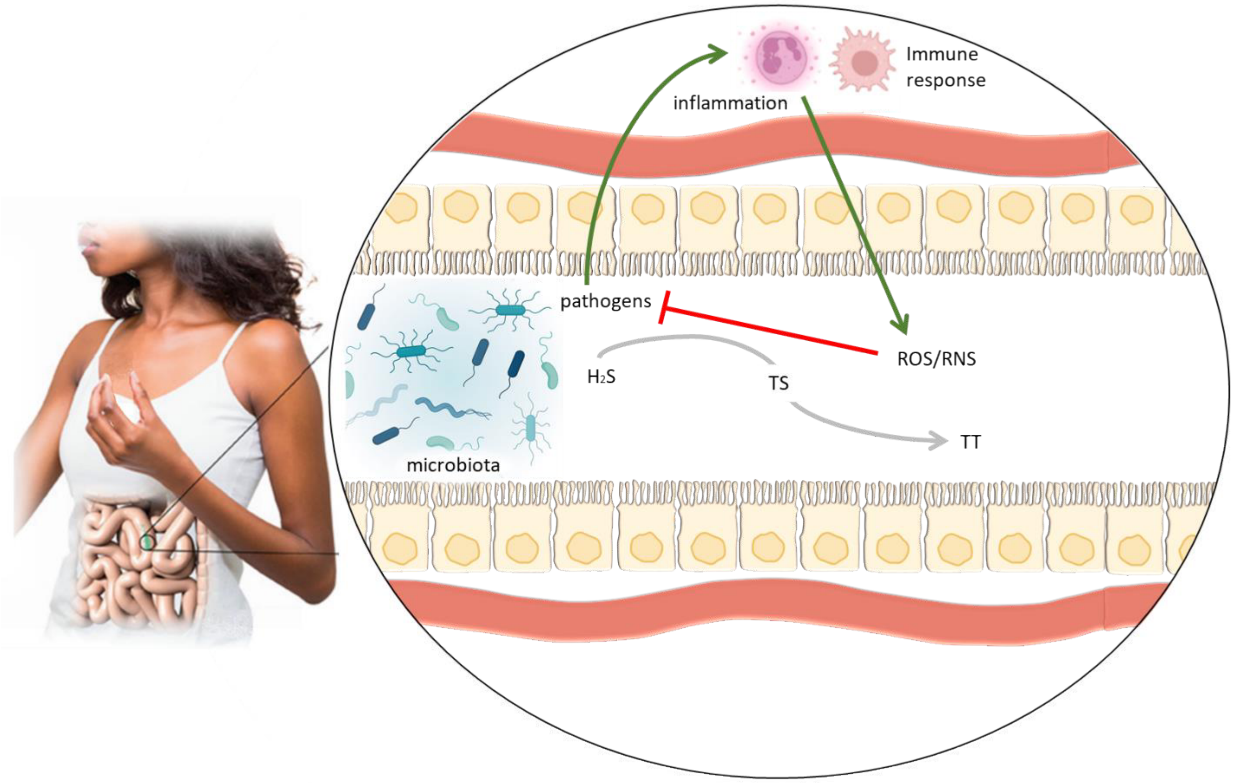


**Fig. S1. Inflammatory bowel disease (IBD) is mediated by labile molecules that are not detectable with current technologies.** Following an inflammatory insult, disproportionate mucosal immune responses via cytokine signaling lead to the release of redox-active molecules such as reactive oxygen species (ROS) and nitric oxide (NO). The resulting oxidative stress inhibits microbial growth in the gut lumen. However, chronic intestinal inflammation damages the epithelium and destroys the epithelial barrier, allowing intestinal microbes to invade the mucosa. The sources of TS in the GI tract are mucin-derived cysteine and sulfate, which are metabolized to $H_2S$[8]. During ulceration, epithelial cells and red blood cells enter the colon; these cells produce enzymes that convert $H_2S$ to TS. In the presence of ROS, TS is oxidized to TT. Consumption of TT and sulfate allows certain pathogens to establish a foot-hold for infection[42,18], evoking further immune responses. These mediators of disease are labile and cannot be measured with existing technology. With only the limited information current approaches provide, breaking this positive feedback loop is challenging. The patient image is reproduced from ref.[50].

A

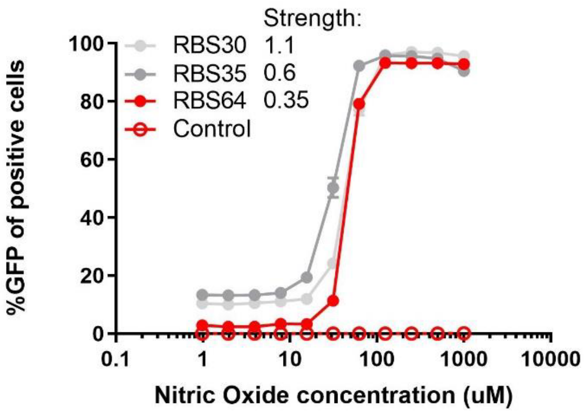
Strength:
RBS30 1.1
RBS35 0.6
RBS64 0.35
Control
%GFP of positive cells
0
20
40
60
80
100
0.1
1
10
100
1000
10000
Nitric Oxide concentration (uM)

B

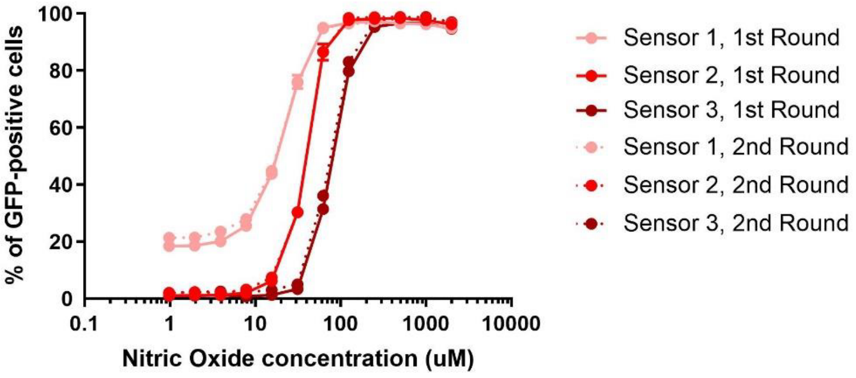
% of GFP-positive cells
0
20
40
60
80
100
0.1
1
10
100
1000
10000
Nitric Oxide concentration (uM)
Sensor 1, 1st Round
Sensor 2, 1st Round
Sensor 3, 1st Round
Sensor 1, 2nd Round
Sensor 2, 2nd Round
Sensor 3, 2nd Round

**C**

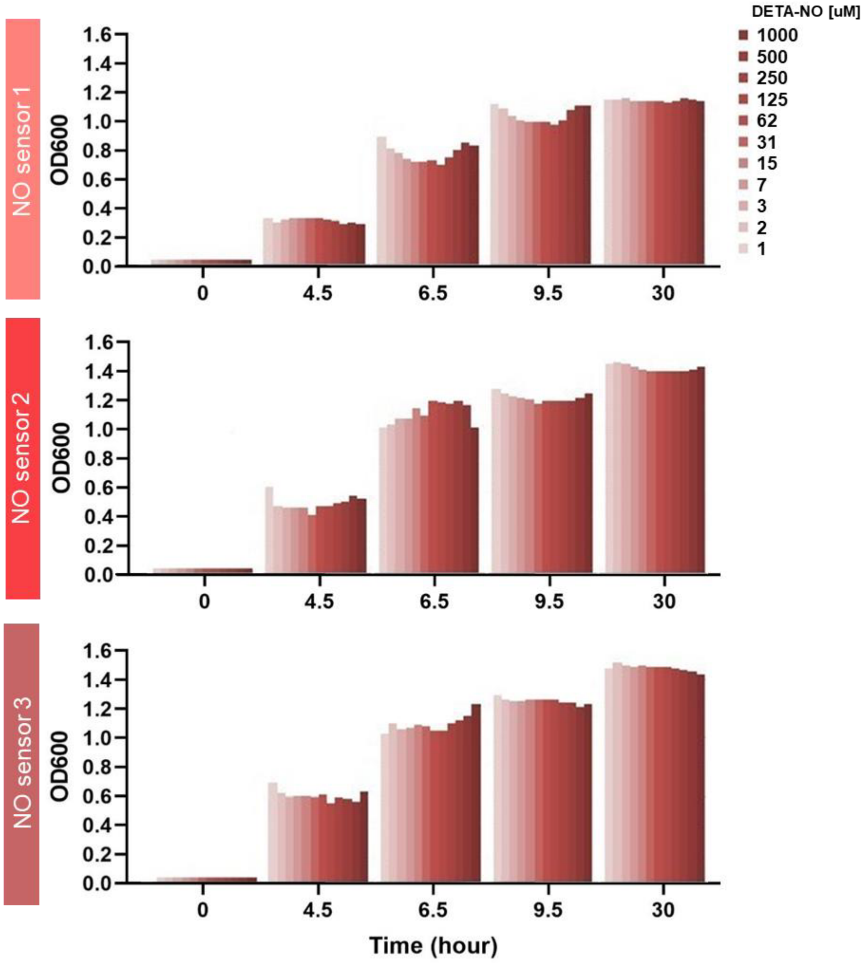

DETA-NO [uM]
1000
500
250
125
62
31
15
7
3
2
1
NO sensor 1
NO sensor 2
NO sensor 3
OD600
1.6
1.4
1.2
1.0
0.8
0.6
0.4
0.2
0.0
0
4.5
6.5
9.5
30
Time (hour)

**D**

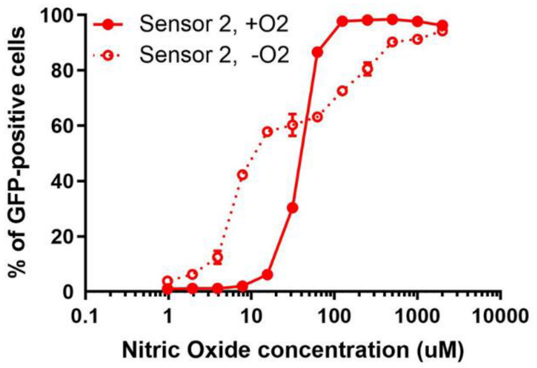


**E**

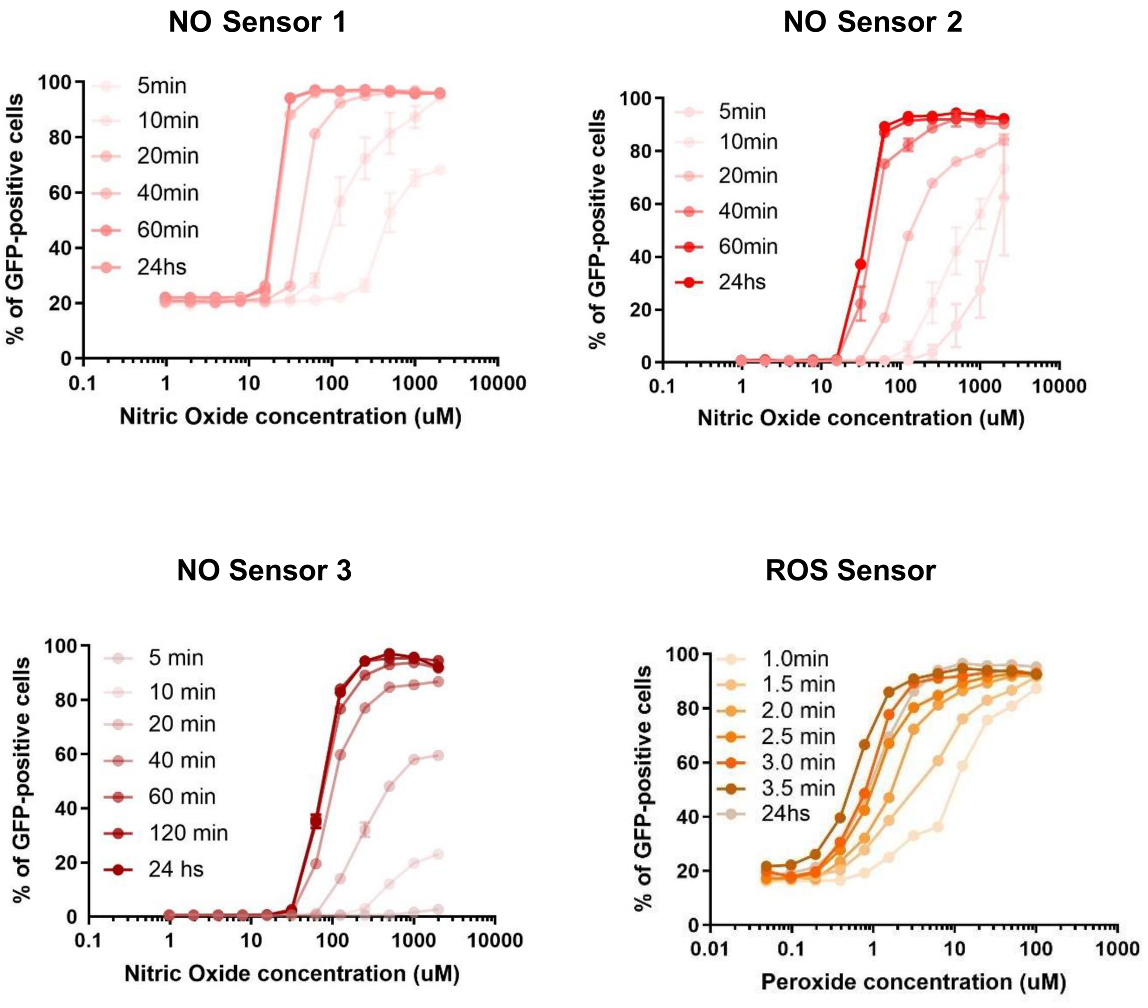


**Fig. S2. A. Genetic circuit optimization and characterization of incorporated recombinase-based switch.** Dose-response curves of NO-sensing genetic circuits in *E. coli* Nissle. The translational initiation strength of the recombianse Bxb1 was varied by using different computationally designed ribosome binding sites (RBS). Predicted RBS strengths are listed in the inset. Lower RBS strength led to a higher SNR. **B.** The memory circuit in the three NO sensors was stable over multiple rounds of re-growth. Engineered bacteria collected in stool were cultured in a selective media to measure NO detection, and the memory system was validated to ascertain that it accurately reflected, over multiple rounds of culturing, the initial input. **C.** GFP expression did not affect growth of bacteria when ON vs. OFF states were compared. **D.** NO detection in anaerobiosis. **E.** Time course of switch activation. The recombinase system triggered GFP expression within minutes (5 mins for sensor NO Sensor 1 and NO Sensor 2, 10 mins for NO Sensor 3 and less than 1 min for the ROS sensor) of exposure to the target molecule. Lines represent the mean. Error bars represent the SEM of two or three independent biological replicates derived from flow cytometry experiments, each of which involved n=10,000 events.

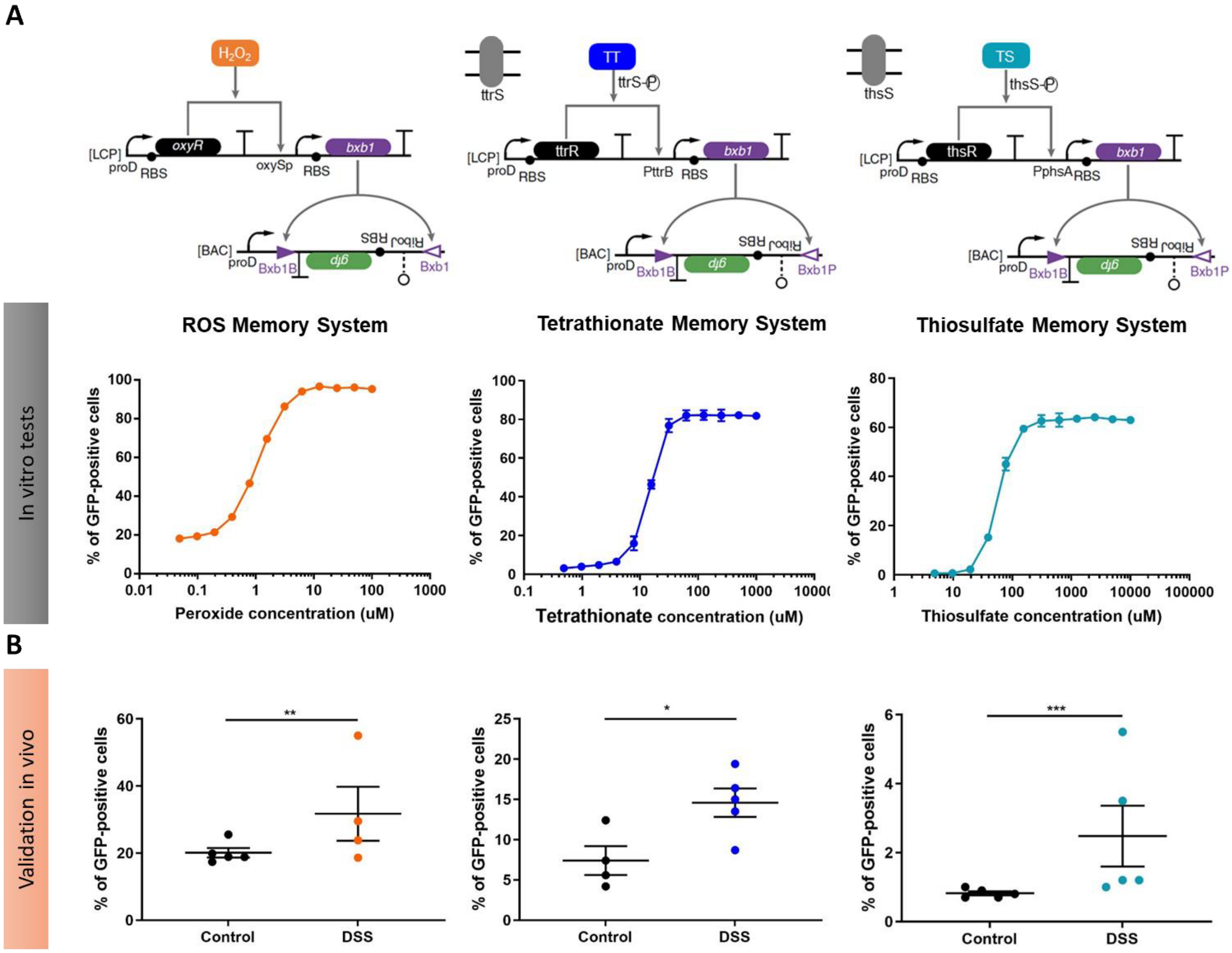

A
H2O2
oxyR
[LCP]
proD
RBS
oxySp
RBS
bxb1
[BAC]
proD
Bxb1B
gfp
RBS
Bxb1
TT
ttrS
ttrS-P
ttrR
PttrB
bxb1
Bxb1P
TS
thsS
thsS-P
thsR
PphsA
ROS Memory System
Tetrathionate Memory System
Thiosulfate Memory System
In vitro tests
% of GFP-positive cells
Peroxide concentration (uM)
Tetrathionate concentration (uM)
Thiosulfate concentration (uM)
B
Validation in vivo
Control
DSS
**
*
***

**C**

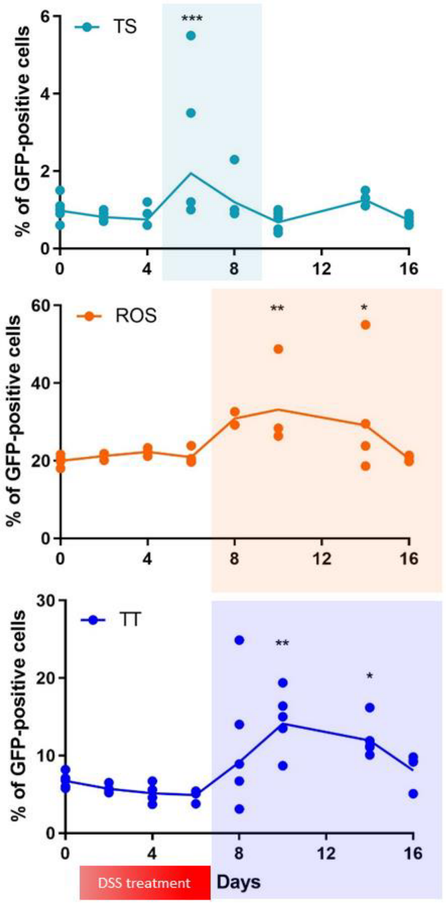


**Fig. S3. Multiple specific disease biomarkers detected in vitro and in vivo. A-C**. The bacterial sensors were validated for the ROS $H_2O_2$, TS, and TT in vitro (A) and in vivo (B-C), following the protocol shown in Fig. 2. In the presence of $H_2O_2$, the transcription factor OxyR is oxidized and activated in *E. coli.* To construct a ROS biosensor that detected $H_2O_2$, the recombinase gene *bxb1* was placed under the control of the OxyR-regulated oxyS promoter, oxySp, on the same genetic circuit[16] (Fig. S3A). To construct the TT and TS sensors, we sought to overcome the oxygen repression that can affect Fumarate and Nitrate Reductase Regulator (FNR)-dependent sensors such as the previously reported two-component system TtrSR[42]. Oxygen levels fluctuate in the gut, depending on the level of disruption of the mucosal epithelium. To avoid this cross-repression, we used two newly identified sensors to express the recombinase system for detecting TS and TT: a TT sensor from *Shewanella baltica*, which does not depend on the FNR system, and the ThsRS sensor from *Shewanella halifaxensis,* the only genetically encoded TS sensor characterized so far[8]. Both sensors distinguished their target molecules from other terminal electron acceptors in vitro[8]. In (A) lines represent the mean, the errors (SEM) are derived from flow cytometry experiments of three representative biological replicates, each of which involved n=10,000 events. In (B-C) individual points represent independent biological replicates, and the bars (days 14, 10 and 6, respectively, for B) and lines

(C) show the mean with SEM. *$p<0.5$, **$p<0.01$, ***$p<0.001$, two-way ANOVA for multiple comparisons.

A

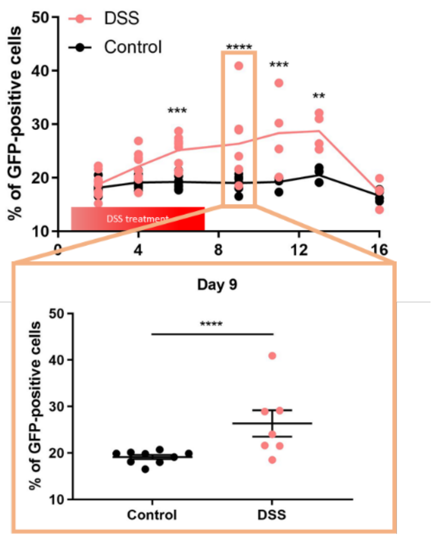
DSS
Control
% of GFP-positive cells
50
40
30
20
10
0
4
8
12
16
***
****
***
**
DSS treatment
Day 9
% of GFP-positive cells
50
40
30
20
10
****
Control
DSS

B

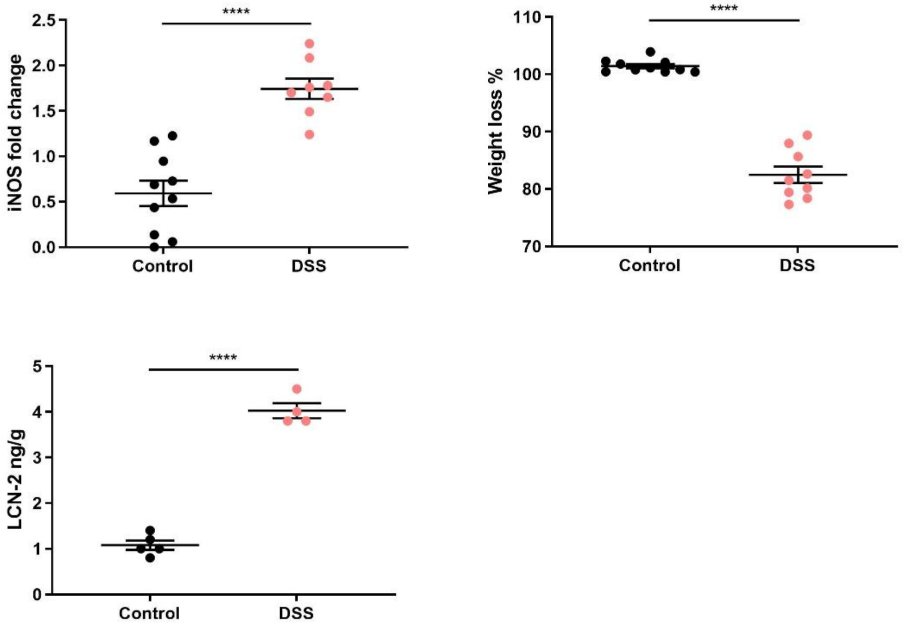
iNOS fold change
2.5
2.0
1.5
1.0
0.5
0.0
****
Control
DSS
Weight loss %
110
100
90
80
70
****
Control
DSS
LCN-2 ng/g
5
4
3
2
1
0
****
Control
DSS

**C**

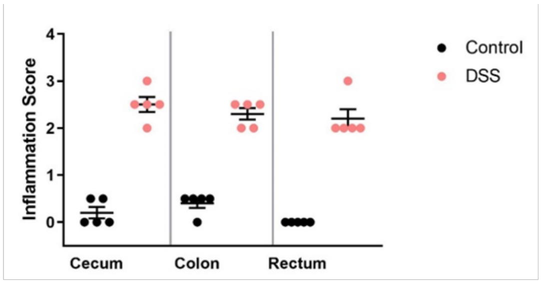


**Inflammation Score**

0 = None

1 = Mild (focal to multifocal low number of inflammatory cells in the lamina propria)

2 = Moderate (multifocal to coalescing moderate number of inflammatory cells restricted to the mucosa and submucosa)

3 = Severe (multifocal to coalescing large number of inflammatory cells affecting all layers of the intestine)

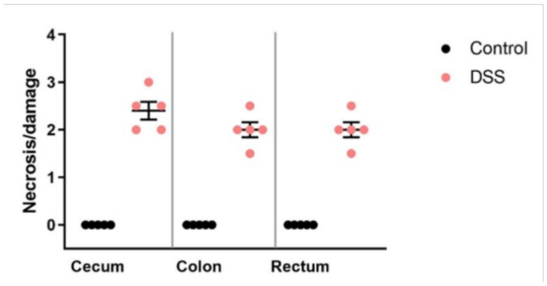


**Necrosis/damage Score**

0 = None

1 = Mild (superficial involving mucosa e.g: superficial ulcerative lesions and erosion)

2 = Moderate (involving mucosa and submucosa, deep ulceration and necrosis, crypt loss, degeneration and atrophy)

3 = Severe (transmural, involving mucosa to serosa)

**D**

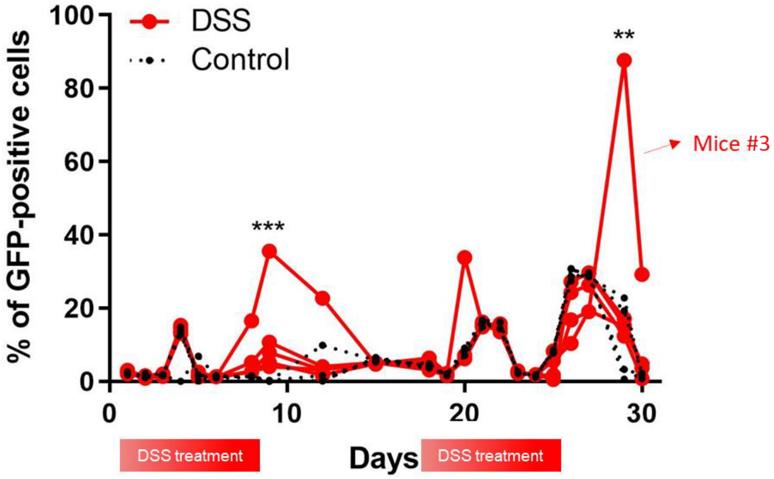

**E**

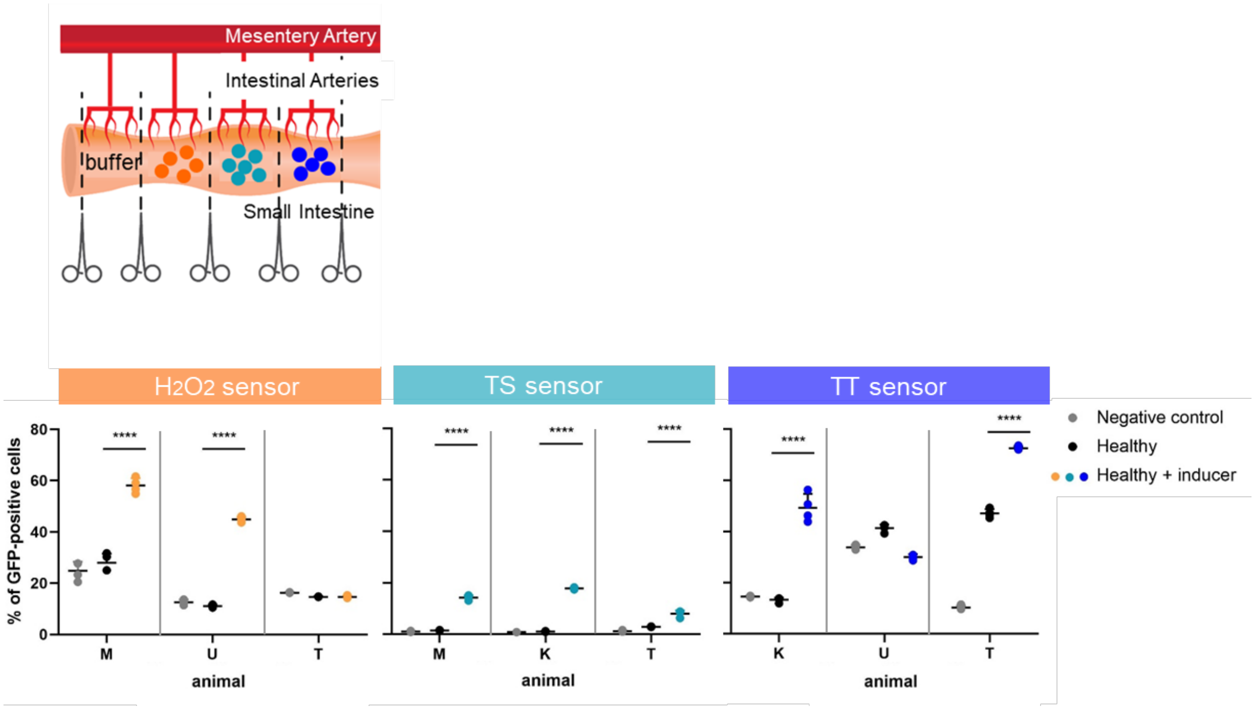


**Fig. S4. In vivo validation of inflammatory biosensors. A.** Detection of NO by the NO biosensor as a marker of GI inflammation in vivo over time. $^{**}p < 0.01$, $^{***}p < 0.001$, $^{****}p < 0.0001$, two-way ANOVA for multiple comparisons. **B.** Independent validation of the presence of inflammation in the DSS colitis model by quantifying iNOS expression during DSS treatment, weight loss, and the lipocalin-2 (LCN-2) biomarker. $^{****}p < 0.0001$, Student's t test. **C.** Histological scores of inflammation and necrosis, indicating the validity of the DSS model. Other indicators were observed but not quantified: bloody and loose stools, poor vigor, anal prolapse, and shortening of the colon upon dissection and gross morphological examination. Lines represent the mean. Error bars represent the SEM of independent biological replicates. **D.** Antibiotic-triggered redox imbalance measured by the NO sensor. NO Sensor 2 allowed us to detect an exacerbated inflammatory response after antibiotic treatment (carbenicillin and chloramphenicol) in a chronic DSS inflammation model, which implies multiple rounds of DSS treatment. Our biosensor for NO shows an increase of NO expression after 4 and 20-30 days of antibiotic treatment in both healthy and DSS-treated mice, with a significant switch activation on day 9 in the DSS-treated mice and on day 29 in the chronic DSS inflammation model, especially high for mouse #3. ''DSS" samples, n = 5 and "Control" samples, n =5. $^{***}p < 0.001$, $^{**}p < 0.01$, two-way ANOVA for multiple comparisons. **E.** Sensor validation in pigs. Experimental design: intestines were clamped to separate the different compartments (control vs. treated), and bacterial sensors were placed in the different compartments (left panel). All sensors registered significant activation in the presence of their respective inducers (300 µM $H_2O_2$, 30 mM TS, 3 mM TT, right panel). The bacteria were collected from the intestine after two hours of exposure to the analyte, and the percent of GFP-positive cells was measured by flow cytometry. Lines represent the mean. The errors (SEM) are derived from flow cytometry experiments of three representative biological replicates, each of which involved n=10,000 events. Here, we show data of three independent experiments (three animals [M, U, T, K] on different days, multiple compartments per animal). **p<0.01, ***p<0.001, ****p<0.0001 Student's t test.

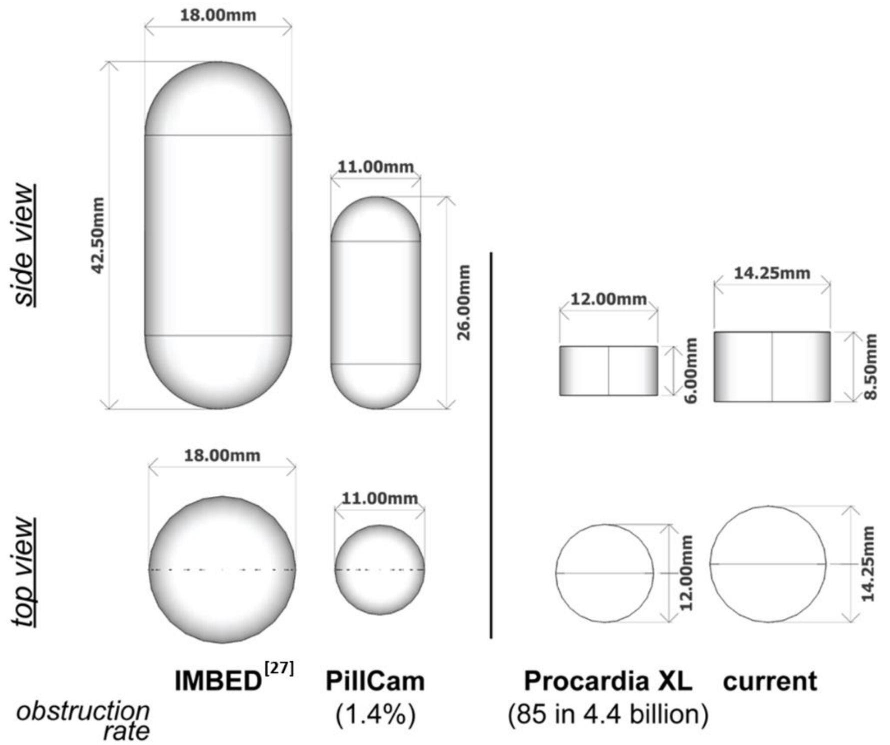


**Fig. S5. Size comparison of ingestible electronics and solid dosage forms with established safety rates.** The safety of ingestible devices depends, in part, on ensuring that these devices will not damage, obstruct, or be retained in the GI tract. The current design was built to conform with the dimensions and form factors of solid dosage forms with known safety profiles and obstruction/retention rates (PillCam, Procardia XL)[46]. Our system integration at the bacterial, electronics and pill casing level allowed a significant reduction in size compared to a previously reported prototype (>9mL to <1.4mL)[27].

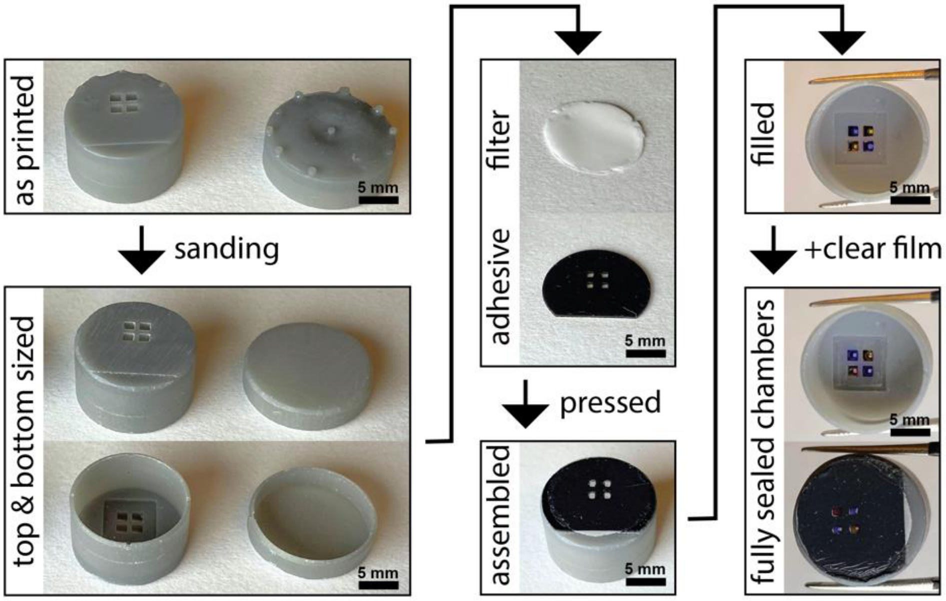


**Fig. S6. Pill casing manufacturing process.** Casing blanks are 3D printed via selective laser sintering (Formlabs) with supports only on the top and bottom face to preserve the thin wall features. The top and bottom faces are then sanded to size. The filter membrane is cut to size with a punch and the double-sided adhesive film is laser cut with through holes aligned to the chambers. After pressing these outer layers together, the chambers are filled with bacterial suspensions from the inside and sealed off with a thin clear adhesive film to yield the fully sealed bacterial chamber/casing unibody. Scale bar = 5 mm.

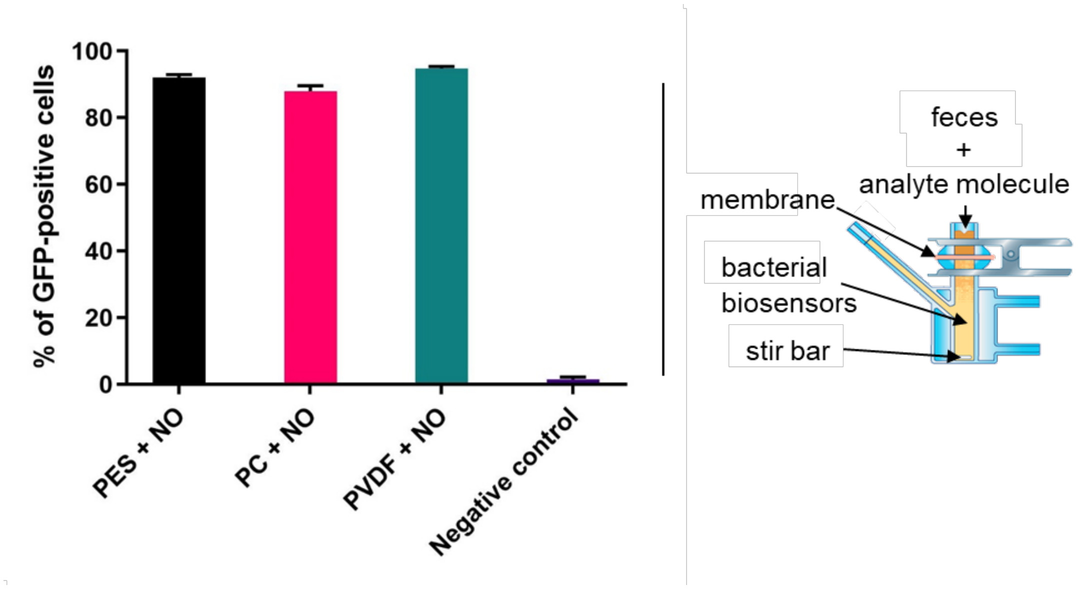


**Fig. S7. Effect of membrane on analyte diffusion.** The porous membranes, when placed in feces, did not interfere with detection of the target molecules. Fouling of the membrane by fecal matter may pose a problem, so we also screened several membrane materials, e.g., polyethersulfone (PES), polycarbonate (PC), and polyvinylidene fluoride (PVDF), to find which material allowed the highest diffusion of the analyte molecules across the membrane. Several porous membranes were tested in Franz Cells (inset on the right) in the presence of feces. All the membranes tested showed similar results, with a similar percentage of detection from the NO bacterial sensors after the analyte had passed through the membrane. The errors (SEM) are derived from flow cytometry experiments of three representative biological replicates, each of which involved n=10,000 events.

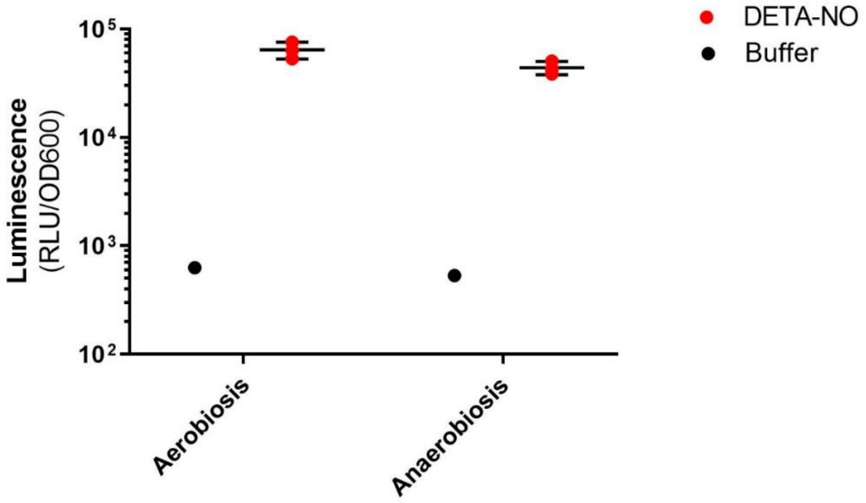


**Fig. S8. Nitric oxide (NO) detection and luciferase expression in anaerobiosis.** Luminescence values were measured overnight post-exposure to the inducer (NO) and normalized to the optical density of the culture. Lines represent the mean. Error bars denote the SEM for three independent biological replicates.

**A**

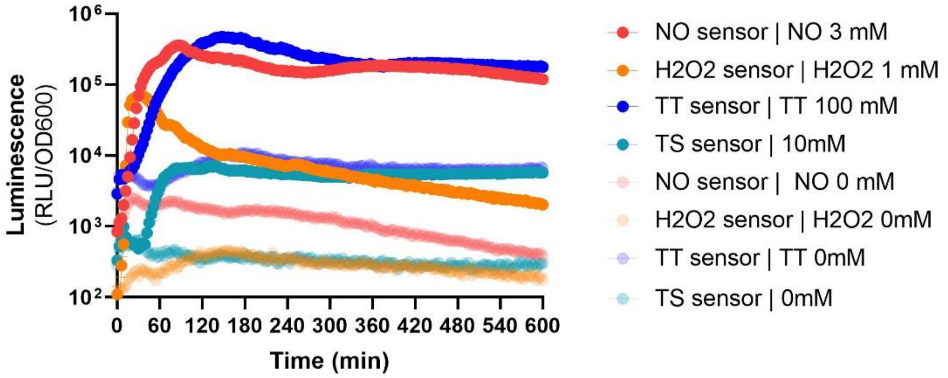


| | SNR |
|---|---|
| NO | 170 |
| H2O2 | 277 |
| TT | 46 |
| TS | 22 |

**B**

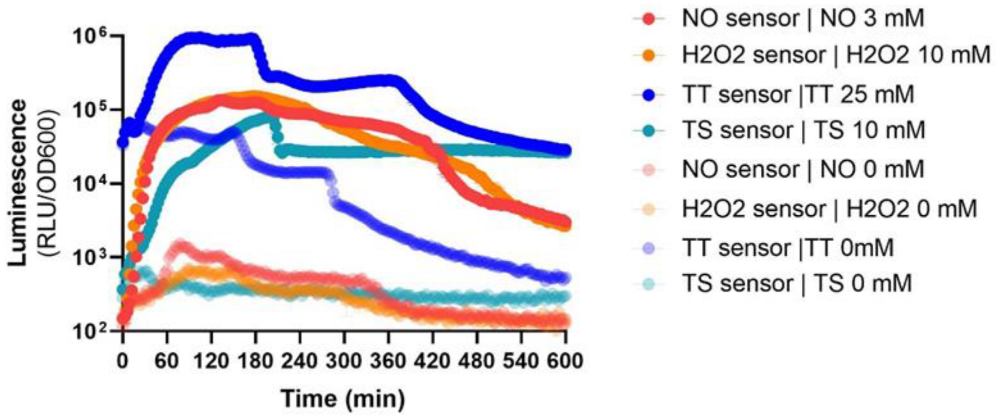


**C**

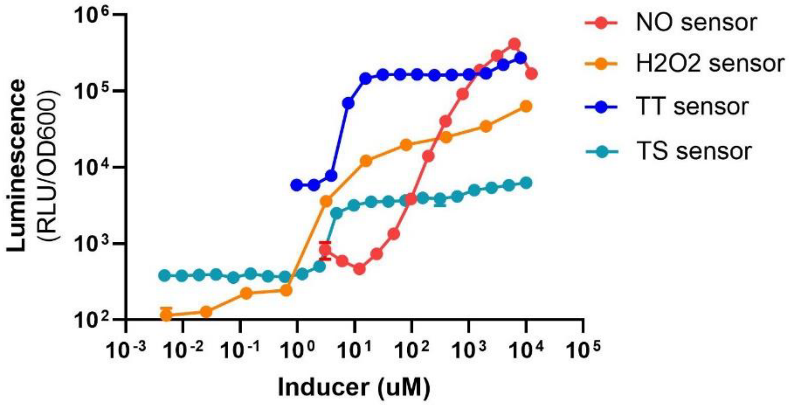


**Fig. S9. Kinetic response of the inflammatory biosensors with the luciferase readout. A-B.** *E. coli* Nissle biosensors were treated with their target analytes in LB (A) or in simulated intestinal fluid (B). The luminescence response was measured in a plate reader every 3 mins for 10 hours. The signal-to-noise ratio (SNR) was calculated by dividing the OD600-normalized luminescence values induced by the OD600-normalized luminescence values of uninduced samples. **C.** Response curve of the inflammatory biosensors with the luciferase readout. *E. coli* Nissle inflammatory sensor strains were treated with various concentrations of their target analytes; maximal luminescence values were measured thirty minutes to two hours post-exposure to the inducer and normalized to the optical density of the culture. Lines represent the mean. Error bars represent the SEM of three independent biological experiments.

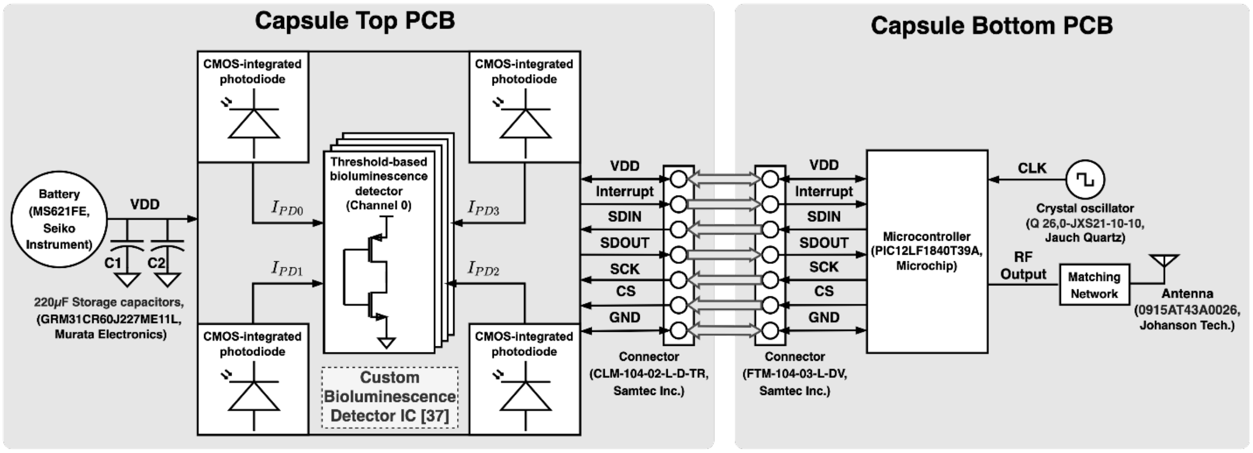


**Fig. S10. Schematic of the ingestible capsule PCB.** The custom bioluminescence detector IC was fabricated in 65 nm CMOS technology[37]. The top PCB holds the custom-designed multi-channel and time-multiplexed bioluminescence detector, a 6.8mm x 2.1mm coin-cell battery, two 220 μF decoupling capacitors, and an 8-position female connector. The bottom PCB holds a microcontroller with an integrated transmitter, a crystal oscillator, an 8-position male connector, an antenna and other components for wireless data transmission. The components on the top and bottom PCB communicate through the two connectors.

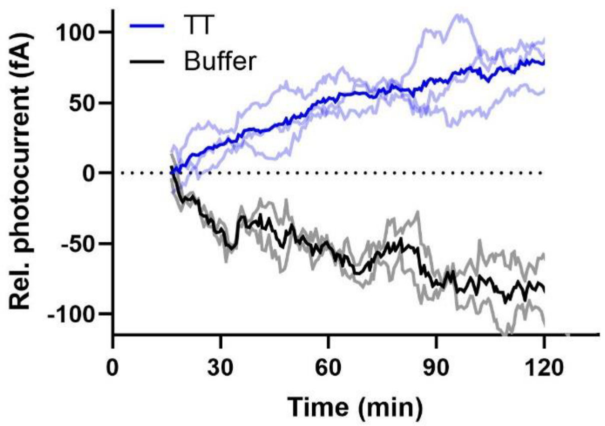


**Fig. S11. Individual replicates of TT sensing in the pig intestinal environment.** The devices with the TT sensor were deposited in the intestinal compartments and TT (100 mM, blue) or buffer alone (black) were injected after temperature stabilization (~15 mins, 37˚C). Readings from the device were wirelessly collected for 120 minutes following device deposition. Dark trace represent the mean of 3 replicate measurements (3 animals on different days, 2 devices per pig, in two different compartments) and pale traces indicate the individual current values for a given device. Photocurrents are provided relative to a one-time calibration value at t=15mins. Non-induced sensor cells (black lines) decrease their luminescence output throughout the experiment (as shown in vitro, tested in simulated intestinal fluid, Fig. S9B), while induced cells express higher levels of luciferase, compensating signal loss over time. For all the replicates, the response of the device placed in the compartment with TT was clearly distinguishable from that of the device in the compartment with the buffer control.

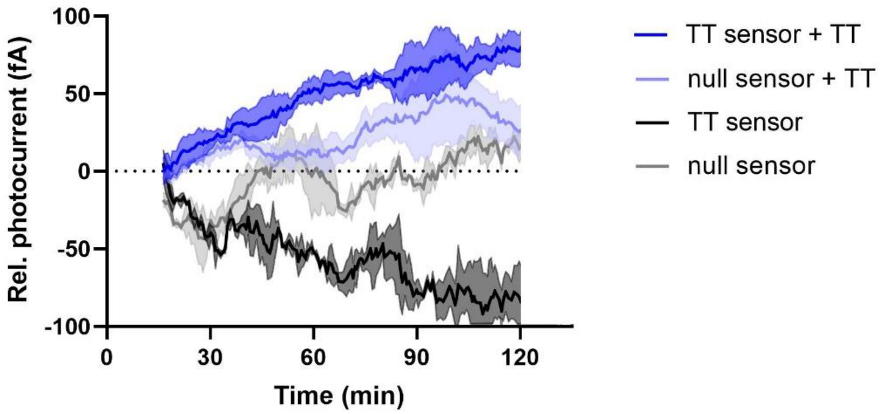


**Fig. S12. Comparison of light detection between different chambers.** *E. coli* Nissle strains containing a functional biosensor circuit for TT detection (TT sensor), and E coli Nissle without the gene for luciferase (null sensor) were loaded into the device. Devices were deposited in the intestine compartments and after temperature stabilization (~15 mins), TT (100 mM, blue) or buffer alone (black) were injected. Compartmentalized intestines were kept inside the abdomen, at 37°C, and wireless signals transmitted from inside the abdomen were collected for 120 mins to analyze the kinetic response of the devices in the abdominal cavity of the pig. Photocurrents provided relative to a one-time calibration value at t=15mins. Non-induced sensor cells (black lines) decrease their luminescence output throughout the experiment (as shown when tested in simulated intestinal fluid, Fig. S9B), while induced cells express higher levels of luciferase compensating signal loss over time. The response of the device placed in the compartment with TT was clearly distinguishable from that of the device in the compartment with the buffer control. Null cells maintain constant values throughout. Error bars denote SEM for three experiments (3 animals on different days, 2 capsules per animal).

### Materials and Methods

#### Bacterial Strains and Culture Conditions

Routine cloning and plasmid propagation were performed in *E. coli* E. cloni 10G (Lucigen). For all in vitro and in vivo experiments, the probiotic strain *E. coli* Nissle 1917 was transformed with gene circuits built on plasmids. Cells were routinely cultured at 37˚C in Luria-Bertani (LB) media (Difco). Where appropriate, growth media was supplemented with antibiotics at the following concentrations: 50 µg/mL kanamycin, 100 µg/mL carbenicillin, 25 µg/mL chloramphenicol, and 100 µg/mL spectinomycin.

#### Plasmid Construction and Circuit characterization

All plasmids were constructed by combining PCR fragments generated by Phusion High-Fidelity DNA Polymerase (NEB) using Gibson Assembly[52], starting from DNA sources as referenced in Table S1 or from gBlocks manufactured by IDT. Genetic parts and plasmids used in this study are listed in Table S1 and will be available from Addgene upon publication. Assembly products were introduced by transformation into chemically competent *E. coli* E. cloni 10G, and sequences were confirmed using Sanger sequencing.

To characterize the constructs built in conjunction with the memory system, appropriate antibiotics were added to Teknova Hi-Def Azure Media containing 0.2% glucose, and *E. coli* E. cloni 10G colonies were inoculated into this culture medium. Cultures were incubated aerobically with shaking for 16–18 hr at 37˚C, then diluted 2,500X into fresh Hi-Def Azure Media (also containing appropriate antibiotics and 0.2% glucose) and incubated aerobically with shaking for another 20 min at 37˚C. Cultures (200 ml) were then transferred to a 96-well plate, and the respective inducers, $H_2O_2$ (hydrogen peroxide Sigma–Aldrich H1009-100ML), DETA/NO (diethylenetriamine/nitric oxide adduct DETA/NO (diethylenetriamine/nitric oxide adduct Sigma–Aldrich D185-50MG), TT (potassium tetrathionate Sigma–Aldrich P2926-100G), and TS (sodium thiosulfate Sigma–Aldrich 217263-250G), were added at appropriate concentrations via serial dilution. Plates were incubated either aerobically with shaking or anaerobically for 20 h at 37˚C for all experiments. For experiments performed in anaerobic conditions, cultures were grown and manipulated in a Coy anaerobic chamber with an atmosphere of 85% $N_2$, 5% $H_2$ and 10% $CO_2$ at 37 °C. All media was pre-reduced overnight in anaerobic atmosphere before inoculation of cultures.

After incubation, the optical densities of cultures were measured at 600 nm in a plate reader. For flow cytometry, cells were diluted in cold 1x PBS and 2% sucrose to reach an optical density of 0.02 (at 600 nm), then assayed on a BD LSRFortessa. The “FITC” channel was used to measure GFP expression. A minimum of 10,000 gated events was recorded. FlowJo software was used to export and process FCS files. Forward scatter and side scatter were used to gate for live cells as described previously[16]. On the resulting flow cytometry histograms, the y-axis is normalized to the mode for each sample. Each experiment was performed for at least three biological replicates.

### Growth and Induction

For genetic circuit characterization, overnight cultures were diluted 1:100 in fresh LB and incubated with shaking at 37˚C for 2 hr. Cultures were removed from the incubator and 200 µL of culture was transferred to a 96-well plate containing various concentrations of inducer. The plate was returned to a shaking incubator at 37˚C. Following 2 hr of incubation, luminescence was read using a BioTek Synergy H1 Hybrid Reader with a 1s integration time and a sensitivity of 150. Luminescence values, measured in relative luminescence units (RLUs), were normalized by the optical density of the culture measured at 600 nm.

For in vitro kinetic studies, subcultured cells were mixed with an inducer in a 96-well plate and immediately placed in the plate reader set at 37˚C without shaking. Luminescence and absorbance were read at 3-minute intervals.

### Mouse Experiments

Approval for mouse experiments was obtained from the Committee on Animal Care at the Massachusetts Institute of Technology. Male C57BL/6J mice (8-10 weeks of age) were obtained from Jackson Labs (Stock No: 000664). Conventional conditions were used to house and handle the mice. One week before the experiments began, the mice were acclimated to the animal facility.

Animals were randomly allocated to experimental groups. Researchers were not blinded to group assignments. Overnight cultures of *E. coli* Nissle grown in Teknova Hi-Def Azure Media with appropriate antibiotics and 0.2% glucose were centrifuged at 5000g for 5 minutes and resuspended in an equal volume of 20% sucrose. Animals were inoculated with 200 µL of bacteria culture (approximately $10^8$ CFU) by oral gavage. Fecal pellets were collected 6 hr post-gavage, and homogenized in 1mL of PBS with a 5 mm stainless steel bead using a TissueLyser II (Qiagen) at 25 Hz for 2 minutes. Samples were centrifuged at 500g for 30 seconds to pellet large fecal debris. Supernatant was cultured in Teknova Hi-Def Azure Media with appropriate antibiotics and 0.2% glucose and incubated aerobically with shaking for 16–18 hr at 37˚C. Cells were then assayed on the flow cytometer.

### Pig Experiments

Approval for pig experiments was obtained from the Committee on Animal Care at the Massachusetts Institute of Technology. Female Yorkshire pigs (50-95kg), received from Cummings Veterinary School at Tufts University in Grafton, MA, were randomly selected for the experiments and housed under conventional conditions. Prior to the experiment, animals were given a clear liquid diet for 24 hours. The day of the experiment, the morning feed was withheld. Pigs were sedated with Telazol® (tiletamine/zolazepam 5 mg/kg), xylazine (2 mg/kg), and atropine (0.04 mg/kg) at the start of the experiment. The jejunum was accessed via a midline laparotomy and the lumen sectioned into several test compartments using Mayo-Robson intestinal clamps. Ischemia-reperfusion injury was used as a model of intestinal inflammation and was caused by clamping the mesentery of the target intestinal segment with hemostatic clamps for 2 hours and then releasing the clamps to allow reperfusion for at least 1 hour. At the end of the procedure pigs were euthanized with Fatal Plus (sodium pentobarbital): 1ml/10lbs or approximately 115~120 mg/kg and heart rate assessed to ensure the pig was euthanized. For bacteria-only experiments, overnight bacterial cultures were diluted 1:10 in LB, cultured for 20 mins, resuspended in 1 mL PBS after centrifugation and were injected into the target intestinal section via a syringe. Healthy intestinal sections were used as is or injected with 200μL of the target analyte (DETA/NO, $H_2O_2$, TT, TS) as described in the text. After the experiment, cells were retrieved by flushing the intestinal section with 10 mL of PBS injected and retrieved via a syringe. For device-bacteria experiments, overnight bacterial cultures were diluted 1:10 in LB, cultured for 20 mins and 1 μL of fresh culture (concentrated 100X by centrifugation) was used to fill the pill casing chambers and sealed as described below (Pill casing manufacture). Devices were inserted into the intestinal lumen through a small incision, manually passed into the target intestinal section and isolated from the incision site by luminal clamping as described above. TT (100 mM) was injected via a syringe into the clamped intestinal compartment and data from the capsules was wirelessly acquired via a 915 MHz radio attached to a laptop. Devices were manually retrieved from the jejunum. A total of 3 animals were included in the experiments; 3 animals on different days, 2 intestinal sections per animal, one for administering the inducer molecule and the other compartment as negative control.

**Preparation of electronic components**

The electronics in the capsules consisted of four photodiodes (Integrated CMOS P+/NWELL/PSUB photodiodes), a custom bioluminescence detector chip fabricated in a CMOS 65 nm process[37], a microcontroller (PIC12LF1840T39A, Microchip) and radio chip (PIC12LF1840T39A, Microchip Technology Inc.), and a 915 MHz chip antenna (0915AT43A0026, Johanson Technology). We used a commercial receiver (CC1200, Texas Instruments). The upper side of the top printed circuit boards (PCB) holds the fully quartz lid-packaged CMOS chip together with an additional on-board LDO (ADP-166, Analog Devices). The assembly was coated with 1 μm of Parylene C to act as a moisture barrier for the electronic components. Parylene C coating was performed using Specialty Coating System Labcoter 2 (PDS 2010) with 1 gram of dichlorodi-p-xylene to reach a target layer thickness of 1 μm using the protocol described by Mimee *et al.* [27].

**Pill casing manufacture**

Pill casing top and bottom blanks were printed via selective laser sintering in Grey resin on a Form 2 printer (Formlabs), post-processed according to the manufacturer's standard protocols and then flat outer faces were sanded to size. In the final design, an Isopore membranes (0.4 μm pore size, Millipore-Sigma) were cut to size with a punch and then attached to the casing body via a thin, laser cut double-sided adhesive layer (3M VHB 5906). Unfilled pill casing tops were conditioned for 24 hours in LB broth, and on the day of the experiment, the chambers were filled from the inside face with bacterial suspensions and subsequently sealed with a thin, laser cut, optically clear adhesive backing film (GeneMate Polyolefin Films with Silicone Adhesive). Bacteria-filled casing tops and empty bottoms were then pressed fit around the electronic system and the outer seam waterproofed with additional silicone (Elite Double 32, Zhermack). Porous membrane types were initially screened as described in Fig. S7 using a standard two-chamber Franz cell.

**In vitro device measurements**

LB culture media was pre-warmed for at least 2 hr prior to the start of experiments. For device-bacterial experiments, overnight cultures were diluted 1:10 in LB, subcultured for 20 min, and concentrated 100X by centrifugation. 1 µL of the concentrated culture was then added to pill casing chambers. Wild-type *E. coli* Nissle 1917 was added to the reference channel for all experiments. Once all four channels were loaded, the cell carrier was fastened to the capsule and fully submerged in pre-warmed media. LB culture media supplemented with inducer (100 mM TT, 20 mM NO, 1 mM $H_2O_2$ or 100 mM TS). Cultures were wrapped several times in thick black fabric to block external light and placed in an incubator at 37˚C, and data was collected wirelessly for 2 hr. At the end of the experiment, devices were disassembled and cell carriers were discarded. Capsules were sterilized with 70% ethanol and thoroughly washed with distilled water. Capsules were left to air-dry and re-used for future experiments.

**Calibration Procedure for Converting Detector Counts to Estimated Photocurrent**

One-time optical calibration was performed to obtain the custom integrated circuit (IC) performance and calibration parameters. During the optical calibration, the wireless capsule holding the custom luminescence detector IC, a standalone photodiode IC, and a green LED ($\lambda = 520\ nm$) were placed inside a metal box covered by a blackout cloth to prevent ambient light from the environment. The capsule and the standalone photodiode IC were placed 1.5 cm adjacent to each other on one side of the metal box, while the LED was placed 30cm opposite to both ICs on the other side of the metal box. We applied five different voltage levels (0 V, 2.1 V, 2.14 V, 2.165 V, 2.185 V) across the LED to obtain different optical power levels. The custom luminescence detector IC wirelessly transmitted sensor readout $N_i$ for each sensing channel $i$ at a given optical power. The standalone photodiode IC was exposed to this optical power simultaneously, and the IC reported a photocurrent level $I_{PD}$ through a sub-femtoamp SourceMeter (K6430, Keithley Instruments). The resolution of a sensing channel $i$ is defined as:

$$Resolution = \frac{\Delta I_{PD}}{\Delta N_i}$$

where $\Delta I_{PD}$ is the difference between photocurrent values reported from the standalone photodiode IC for two LED bias voltages; and $\Delta N_i$ is the difference between luminescence detector IC output counts for the same LED bias voltages. The resolution for a single channel was calculated using a linear regression model to fit the luminescence IC output counts and photocurrents from standalone photodiode IC over five LED bias voltages. The resolution of the luminescence detector chip is 5.8-6.5 fA/count. The minimum detectable signal at one LED bias voltage was calculated using the resolution of each sensing channel times $1\sigma$ standard deviation of the IC output at this optical power. The worst-case minimum detectable signal is 71 fA. The estimated photocurrents for in-vitro and in-vivo measurements were calculated by taking the product of resolutions from one-time optical calibration and measured IC output counts in each measurement. We used a moving average filter of 25 samples (~15 min moving average).

**Data Analysis, Statistics and Computational Methods**

All data were analyzed using GraphPad Prism version 9.1.2 (Graph Software, San Diego, CA, USA, http://www.graphpad.com). Sequences were analyzed using SnapGene version 5.1.2

(www.snapgene.com). As noted, error bars represent the standard error of the mean (SEM) of at least three independent experiments carried out on different days. Significant differences between groups was determined using an unpaired, two-tailed Student's t-test assuming unequal variance and for curves over time, two-way ANOVA for multiple comparisons. Fold change or signal-to-noise ratio was determined by dividing the normalized luminescence values (RLU/OD600) of samples treated with the maximal inducer concentration with uninduced samples. The receiver operating characteristic was calculated based on three independent experiments. Fig. 1 and Fig. S1 were created with BioRender.com.
