## Supplementary material for "Ingestible capsule for detecting labile inflammatory biomarkers in situ": Table S1

**Table S1. Genetic Parts**

**Part Name**. DNA sequence

**oxySp**

TTCATTATCCATCCTCCATCGCCACGATAGTTCATGGCGATAGGTAGAATAGCAA
TGAACGATTATCCCTATCAAGCATTCTGACTGAGCATTGCTCACA

**oxyR**

ATGAATATTCGTGATCTTGAGTACCTGGTGGCATTGGCTGAACACCGCCATTTT
CGGCGTGCGGCAGATTCCTGCCACGTTAGCCAGCCGACGCTTAGCGGGCAAATTCGT
AAGCTGGAAGATGAGCTGGGCGTGATGTTGCTGGAGCGGACCAGCCGTAAAGTGTTGT
TCACCCAGGCGGGAATGCTGCTGGTGGATCAGGCGCGTACCGTGCTGCGTGAGGTGA
AAGTCCTTAAAGAGATGGCAAGCCAGCAGGGCGAGACGATGTCCGGACCGCTGCACA
TTGGTTTGATTCCCACAGTTGGACCGTACCTGCTACCGCATATTATCCCTATGCTGCAC
CAGACCTTTCCAAAGCTGGAAATGTATCTGCATGAAGCACAGACCCACCAGTTACTGGC
GCAACTGGACAGCGGCAAACTCGATTGCGTGATCCTCGCGCTGGTGAAAGAGAGCGA
AGCATTCATTGAAGTGCCGTTGTTTGATGAGCCAATGTTGCTGGCTATCTATGAAGATC
ACCCGTGGGCGAACCGCGAATGCGTACCGATGGCCGATCTGGCAGGGGAAAAACTGC
TGATGCTGGAAGATGGTCACTGTTTGCGCGATCAGGCAATGGGTTTCTGTTTTGAAGCC
GGGGCGGATGAAGATACACACTTCCGCGCGACCAGCCTGGAAACTCTGCGCAACATG
GTGGCGGCAGGTAGCGGGATCACTTTACTGCCAGCGCTGGCTGTGCCGCCGGAGCGC
AAACGCGATGGGGTTGTTTATCTGCCGTGCATTAAGCCGGAACCACGCCGCACTATTG
GCCTGGTTTATCGTCCTGGCTCACCGCTGCGCAGCCGCTATGAGCAGCTGGCAGAGG
CCATCCGCGCAAGAATGGATGGCCATTTCGATAAAGTTTTAAAACAGGCGGTTTAA

**gfp**

ATGAGTAAAGGAGAAGAACTTTTCACTGGAGTTGTCCCAATTCTTGTTGAATTAG
ATGGTGATGTTAATGGGCACAAATTTTCTGTCAGTGGAGAGGGTGAAGGTGATGCAACA
TACGGAAAACTTACCCTTAAATTTATTTGCACTACTGGAAAACTACCTGTTCCATGGCCA
ACACTTGTCACTACTTTCGGTTATGGTGTTCAATGCTTTGCGAGATACCCAGATCATATG
AAACAGCATGACTTTTTCAAGAGTGCCATGCCCGAAGGTTATGTACAGGAAAGAACTAT
ATTTTTCAAAGATGACGGGAACTACAAGACACGTGCTGAAGTCAAGTTTGAAGGTGATA
CCCTTGTTAATAGAATCGAGTTAAAAGGTATTGATTTTAAAGAAGATGGAAACATTCTTG
GACACAAATTGGAATACAACTATAACTCACACAATGTATACATCATGGCAGACAAACAAA
AGAATGGAATCAAAGTTAACTTCAAAATTAGACACAACATTGAAGATGGAAGCGTTCAAC
TAGCAGACCATTATCAACAAAATACTCCAATTGGCGATGGCCCTGTCCTTTTACCAGAC
AACCATTACCTGTCCACACAATCTGCCCTTTCGAAAGATCCCAACGAAAAGAGAGACCA
CATGGTCCTTCTTGAGTTTGTAACAGCTGCTGGGATTACACATGGCATGGATGATCTCT
ACAAATAA

**Bxb1**

ATGAGAGCCCTGGTAGTCATCCGCCTGTCCCGCGTCACCGATGCTACGACTTC
ACCGGAGCGTCAGCTGGAGTCTTGCCAGCAGCTCTGCGCCCAGCGCGGCTGGGACGT
CGTCGGGGTAGCGGAGGATCTGGACGTCTCCGGGGCGGTCGATCCGTTCGACCGGAA
GCGCAGACCGAACCTGGCCCGGTGGCTAGCGTTCGAGGAGCAACCGTTCGACGTGAT
CGTGGCGTACCGGGTAGACCGGTTGACCCGATCGATCCGGCATCTGCAGCAGCTGGT
CCACTGGGCCGAGGACCACAAGAAGCTGGTCGTCTCCGCGACCGAAGCGCACTTCGA
TACGACGACGCCGTTTGCGGCGGTCGTCATCGCGCTTATGGGAACGGTGGCGCAGAT
GGAATTAGAAGCGATCAAAGAGCGGAACCGTTCGGCTGCGCATTTCAATATCCGCGCC
GGGAAATACCGAGGATCCCTGCCGCCGTGGGGATACCTGCCTACGCGCGTGGACGGG
GAGTGGCGGCTGGTGCCGGACCCTGTGCAGCGAGAGCGCATCCTCGAGGTGTATCAC
CGCGTCGTCGACAACCACGAGCCGCTGCACCTGGTGGCCCACGACCTGAACCGGCGT

GGTGTCCTGTCGCCGAAGGACTACTTCGCGCAGCTGCAAGGCCGCGAGCCGCAGGGC
CGGGAGTGGTCGGCTACCGCGCTGAAGCGATCGATGATCTCCGAGGCGATGCTCGGG
TACGCGACTCTGAACGGTAAGACCGTCCGAGACGACGACGGAGCCCCGCTGGTGCGG
GCTGAGCCGATCCTGACCCGTGAGCAGCTGGAGGCGCTGCGCGCCGAGCTCGTGAAG
ACCTCCCGGGCGAAGCCCGCGGTGTCTACCCCGTCGCTGCTGCTGCGGGTGTTGTTC
TGTGCGGTGTGCGGGGAGCCCGCGTACAAGTTCGCCGGGGGAGGACGTAAGCACCC
GCGCTACCGCTGCCGCTCGATGGGGTTCCCGAAGCACTGCGGGAACGGCACGGTGG
CGATGGCCGAGTGGGACGCGTTCTGCGAGGAGCAGGTGCTGGATCTGCTCGGGGAC
GCGGAGCGTCTGGAGAAAGTCTGGGTAGCCGGCTCGGACTCCGCGGTCGAACTCGCG
GAGGTGAACGCGGAGCTGGTGGACCTGACGTCGCTGATCGGCTCCCCGGCCTACCGG
GCCGGCTCTCCGCAGCGAGAAGCACTGGATGCCCGTATTGCGGCGCTGGCCGCGCG
GCAAGAGGAGCTGGAGGGTCTAGAGGCTCGCCCGTCTGGCTGGGAGTGGCGCGAGA
CCGGGCAGCGGTTCGGGGACTGGTGGCGGGAGCAGGACACCGCGGCAAAGAACACC
TGGCTTCGGTCGATGAACGTTCGGCTGACGTTCGACGTCCGCGGCGGGCTGACTCGC
ACGATCGACTTCGGGGATCTGCAGGAGTACGAGCAGCATCTCAGGCTCGGCAGCGTG
GTCGAACGGCTACACACCGGGATGTCGAGGCCTGCAGCAAACGACGAAAACTACGCT
GCAGCAGTTTAG

**J23104**

TTGACAGCTAGCTCAGTCCTAGGTATTGTGCTAGCCTAGTATCGATCTCCATAAC
TATCCTATAGATC

**J23105**

TTTACGGCTAGCTCAGTCCTAGGTACTATGCTAGCAGAAATATAAAGAACGATCT
ATTTATCCGCGTAC

**ThsS**

ATGTCCCGCCTGCTGCTGTGTATCTGTGTTCTGCTGTTCTCTTCTGTGGCGTGG
TCTAAACCGCAGCAGTTTTATGTGGGCGTACTGGCTAACTGGGGTCATCAGCAAGCCG
TTGAACGTTGGACCCCGATGATGGAGTATCTGAACGAACATGTGCCGGACGCGGAATT
TCACGTCTACCCGGGCAACTTCAAAGCACTGAACCTGGCAATGGAACTGGGCCAGATT
CAGTTCATTATCACTAACCCGGGCCAATATCTGTACCTGAGCAATCAGTACCCGCTGTC
TTGGCTGGCGACCATGCGTTCTAAGCGTCACGATGGTACCACTTCTGCGATCGGTTCC
GCCATTATTGTCCGCGCGGACAGCGACTACCGCACCCTGTACGACCTGAAAGGTAAAG
TGGTGGCTGCGTCCGACCCGCATGCTCTGGGTGGCTACCAAGCGACCGTCGGTCTGA
TGCATTCCCTGGGCATGGATCCGGACACCTTCTTCGGTGAAACCAAGTTTCTGGGCTTT
CCACTGGATCCGCTGCTGTACCAAGTTCGTGATGGCAACGTTGACGCGGCCATTACCC
CACTGTGCACTCTGGAGGACATGGTTGCACGCGGCGTACTGAAATCTTCCGATTTTCGT
GTGCTGAACCCTAGCCGCCCGGATGGTGTAGAATGCCAGTGCTCTACCACCCTGTACC
CGAACTGGTCTTTCGCTGCGACTGAGTCTGTATCCACCGAACTGTCTAAAGAAATCACG
CAGGCACTGCTGGAACTGCCATCCGACAGCCCGGCAGCTATCAAAGCGCAACTGACC
GGCTGGACCAGCCCGATCTCCCAACTGGCGGTAATCAAACTGTTCAAAGAGCTGCACG
TAAAAACCCCGGACTCTAGCCGTTGGGAAGCCGTTAAGAAGTGGCTGGAAGAAAACCG
TCACTGGGGTATCCTGTCTGTTCTGGTGTTCATCATTGCAACGCTGTATCACCTGTGGA
TTGAATACCGCTTCCACCAAAAAAGCTCTTCTCTGATCGAATCTGAACGTCAGCTGAAA
CAGCAAGCTGTTGCCCTGGAACGTCTGCAATCTGCTAGCATCGTTGGTGAAATTGGTG
CGGGTCTGGCCCACGAGATTAATCAGCCGATCGCTGCAATTACCTCTTATTCTGAAGGT
GGCATCATGCGCCTGCAAGGTAAAGAACAGGCGGATACGGATAGCTGCATCGAACTGC
TGGAAAAAATCCACAAACAGAGCACTCGCGCAGGCGAAGTGGTGCACCGCATCCGTG
GTCTGCTGAAACGTCGTGAAGCGGTGATGGTAGATGTTAACATCCTGACCCTGGTGGA
AGAATCCATCAGCCTGCTGCGTCTGGAGCTGGCACGTCGCGAAATCCAGATCAACACT
CAGATCAAAGGTGAACCGTTCTTCATTACTGCCGACCGCGTTGGCCTGCTGCAAGTTCT

GATTAACCTGATCAAAAACTCCCTGGACGCGATCGCTGAATCTGATAATGCCCGTTCTG
GTAAAATCAACATCGAACTGGACTTTAAAGAGTACCAGGTAAACGTCTCCATCATCGAT
AACGGTCCGGGCCTGGCGATGGATTCTGACACTCTGATGGCTACGTTTTACACTACCA
AAATGGATGGCCTGGGCCTGGGTCTGGCAATCTGCCGCGAAGTTATCAGCAACCACGA
CGGCCACTTCCTGCTGTCCAACCGTGACGACGGCGTTCTGGGCTGTGTGGCAACCCT
GAATCTGAAAAAACGCGGTTCTGAAGTGCCGATCGAAGTCTAA

**ThsR**

ATGCAGCAGCAAATCAACGGCCCGGTCTACCTGGTGGATGATGATGAAGCCAT
TATCGACTCCATCGATTTTTTGATGGAGGGCTACGGTTACAAACTGAACTCGTTTAACTG
CGGCGATCGCTTTTTGGCAGAAGTCGATCTCACCCAGGCAGGATGTGTAATTCTGGAT
GCGCGTATGCCAGGCTTAACTGGTCCTCAGGTGCAACAGCTGCTGAGCGACGCGAAAA
GCCCGCTTGCGGTCATCTTCCTGACCGGCCATGGCGATGTTCCGATGGCGGTTGATGC
GTTCAAAAATGGCGCGTTCGATTTCTTTCAAAAACCTGTGCCGGGTAGCTTGCTCAGTC
AGTCAATTGCCAAAGGCTTGACTTATTCAATCGATCAACATCTGAAACGTACTAACCAAG
CGTTAATCGACACGCTCTCGGAACGCGAAGCTCAAATTTTTCAACTGGTGATTGCAGGC
AACACCAACAAACAGATGGCTAACGAGCTTTGCGTGGCTATTCGTACCATTGAGGTTCA
CCGTAGCAAACTGATGACCAAACTGGGTGTTAACAACCTGGCTGAACTGGTTAAACTGG
CGCCGCTGCTGGCACATAAATCCGAATAA

**PphsA**

TTCAAGCATTATTATGCTGTTTTTTGAAGTGAATGTGCGGCCATCTAGCCGCACA
TTTTGCATCTAAAACATGCAGTCATCAGCAAAATAATAAACTTTTCCCCAATATGTGGTTT
ACCACAATTTACAGGAATTCACTCCTGTGGTGGTGCAAATTTGAACTGTGAATTGCTTCA
CAAACGCCGCTATCGCAATGTCAGTATGTGGTTTACCACAATATCTAATATCACTCTGCT
CAATAACAATGATGAAAACCTTAGGAAGAAGTTAATTGTGTTAAACAGTTAACTAGGGGC
TTTATCTAACGCTCTCCTAAGGACAACTGTCATTGGGAGATTTAAC

**LuxCDABE**

ATGACTAAAAAAATTTCATTCATTATTAACGGCCAGGTTGAAATCTTTCCCGAAA
GTGATGATTTAGTGCAATCCATTAATTTTGGTGATAATAGTGTTTACCTGCCAATATTGA
ATGACTCTCATGTAAAAAACATTATTGATTGTAATGGAAATAACGAATTACGGTTGCATA
ACATTGTCAATTTTCTCTATACGGTAGGGCAAAGATGGAAAAATGAAGAATACTCAAGAC
GCAGGACATACATTCGTGACTTAAAAAAATATATGGGATATTCAGAAGAAATGGCTAAG
CTAGAGGCCAATTGGATATCTATGATTTTATGTTCTAAAGGCGGCCTTTATGATGTTGTA
GAAAATGAACTTGGTTCTCGCCATATCATGGATGAATGGCTACCTCAGGATGAAAGTTA
TGTTCGGGCTTTTCCGAAAGGTAAATCTGTACATCTGTTGGCAGGTAATGTTCCATTATC
TGGGATCATGTCTATATTACGCGCAATTTTAACTAAGAATCAGTGTATTATAAAAACATC
GTCAACCGATCCTTTTACCGCTAATGCATTAGCGTTAAGTTTTATTGATGTAGACCCTAA
TCATCCGATAACGCGCTCTTTATCTGTTATATATTGGCCCCACCAAGGTGATACATCACT
CGCAAAAGAAATTATGCGACATGCGGATGTTATTGTCGCTTGGGGAGGGCCAGATGCG
ATTAATTGGGCGGTAGAGCATGCGCCATCTTATGCTGATGTGATTAAATTTGGTTCTAAA
AAGAGTCTTTGCATTATCGATAATCCTGTTGATTTGACGTCCGCAGCGACAGGTGCGGC
TCATGATGTTTGTTTTTACGATCAGCGAGCTTGTTTTTCTGCCCAAAACATATATTACATG
GGAAATCATTATGAGGAATTTAAGTTAGCGTTGATAGAAAAACTTAATCTATATGCGCAT
ATATTACCGAATGCCAAAAAAGATTTTGATGAAAAGGCGGCCTATTCTTTAGTTCAAAAA
GAAAGCTTGTTTGCTGGATTAAAAGTAGAGGTGGATATTCATCAACGTTGGATGATTATT
GAGTCAAATGCAGGTGTGGAATTTAATCAACCACTTGGCAGATGTGTGTACCTTCATCA
CGTCGATAATATTGAGCAAATATTGCCTTATGTTCAAAAAAATAAGACGCAAACCATATC
TATTTTTCCTTGGGAGTCATCATTTAAATATCGAGATGCGTTAGCATTAAAAGGTGCGGA
AAGGATTGTAGAAGCAGGAATGAATAACATATTTCGAGTTGGTGGATCTCATGACGGAA
TGCGACCGTTGCAACGATTAGTGACATATATTTCTCATGAAAGGCCATCTAACTATACG

GCTAAGGATGTTGCGGTTGAAATAGAACAGACTCGATTCCTGGAAGAAGATAAGTTCCT
TGTATTTGTCCCATAATAGGTAAAAGTATGGAAAATGAATCAAAATATAAAACCATCGAC
CACGTTATTTGTGTTGAAGGAAATAAAAAAATTCATGTTTGGGAAACGCTGCCAGAAGA
AAACAGCCCAAAGAGAAAGAATGCCATTATTATTGCGTCTGGTTTTGCCCGCAGGATGG
ATCATTTTGCTGGTCTGGCGGAATATTTATCGCGGAATGGATTTCATGTGATCCGCTAT
GATTCGCTTCACCACGTTGGATTGAGTTCAGGGACAATTGATGAATTTACAATGTCTATA
GGAAAGCAGAGCTTGTTAGCAGTGGTTGATTGGTTAACTACACGAAAAATAAATAACTT
CGGTATGTTGGCTTCAAGCTTATCTGCGCGGATAGCTTATGCAAGCCTATCTGAAATCA
ATGCTTCGTTTTTAATCACCGCAGTCGGTGTTGTTAACTTAAGATATTCTCTTGAAAGAG
CTTTAGGGTTTGATTATCTCAGTCTACCCATTAATGAATTGCCGGATAATCTAGATTTTG
AAGGCCATAAATTGGGTGCTGAAGTCTTTGCGAGAGATTGTCTTGATTTTGGTTGGGAA
GATTTAGCTTCTACAATTAATAACATGATGTATCTTGATATACCGTTTATTGCTTTTACTG
CAAATAACGATAATTGGGTCAAGCAAGATGAAGTTATCACATTGTTATCAAATATTCGTA
GTAATCGATGCAAGATATATTCTTTGTTAGGAAGTTCGCATGACTTGAGTGAAAATTTAG
TGGTCCTGCGCAATTTTTATCAATCGGTTACGAAAGCCGCTATCGCGATGGATAATGAT
CATCTGGATATTGATGTTGATATTACTGAACCGTCATTTGAACATTTAACTATTGCGACA
GTCAATGAACGCCGAATGAGAATTGAGATTGAAAATCAAGCAATTTCTCTGTCTTAAAAT
CTATTGAGATATTCTATCACTCAAATAGCAATATAAGGACTCTCTATGAAATTTGGAAACT
TTTTGCTTACATACCAACCTCCCCAATTTTCTCAAACAGAGGTAATGAAACGTTTGGTTA
AATTAGGTCGCATCTCTGAGGAGTGTGGTTTTGATACCGTATGGTTACTGGAGCATCAT
TTCACGGAGTTTGGTTTGCTTGGTAACCCTTATGTCGCTGCTGCATATTTACTTGGCGC
GACTAAAAAATTGAATGTAGGAACTGCCGCTATTGTTCTTCCCACAGCCCATCCAGTAC
GCCAACTTGAAGATGTGAATTTATTGGATCAAATGTCAAAAGGACGATTTCGGTTTGGTA
TTTGCCGAGGGCTTTACAACAAGGACTTTCGCGTATTCGGCACAGATATGAATAACAGT
CGCGCCTTAGCGGAATGCTGGTACGGGCTGATAAAGAATGGCATGACAGAGGGATATA
TGGAAGCTGATAATGAACATATCAAGTTCCATAAGGTAAAAGTAAACCCCGCGGCGTAT
AGCAGAGGTGGCGCACCGGTTTATGTGGTGGCTGAATCAGCTTCGACGACTGAGTGG
GCTGCTCAATTTGGCCTACCGATGATATTAAGTTGGATTATAAATACTAACGAAAAGAAA
GCACAACTTGAGCTTTATAATGAAGTGGCTCAAGAATATGGGCACGATATTCATAATATC
GACCATTGCTTATCATATATAACATCTGTAGATCATGACTCAATTAAAGCGAAAGAGATT
TGCCGGAAATTTCTGGGGCATTGGTATGATTCTTATGTGAATGCTACGACTATTTTTGAT
GATTCAGACCAAACAAGAGGTTATGATTTCAATAAAGGGCAGTGGCGTGACTTTGTATT
AAAAGGACATAAAGATACTAATCGCCGTATTGATTACAGTTACGAAATCAATCCCGTGG
GAACGCCGCAGGAATGTATTGACATAATTCAAAAAGACATTGATGCTACAGGAATATCA
AATATTTGTTGTGGATTTGAAGCTAATGGAACAGTAGACGAAATTATTGCTTCCATGAAG
CTCTTCCAGTCTGATGTCATGCCATTTCTTAAAGAAAAACAACGTTCGCTATTATATTAG
CTAAGGAGAAAGAAATGAAATTTGGATTGTTCTTCCTTAACTTCATCAATTCAACAACTG
TTCAAGAACAAAGTATAGTTCGCATGCAGGAAATAACGGAGTATGTTGATAAGTTGAATT
TTGAACAGATTTTAGTGTATGAAAATCATTTTTCAGATAATGGTGTTGTCGGCGCTCCTC
TGACTGTTTCTGGTTTTCTGCTCGGTTTAACAGAGAAAATTAAAATTGGTTCATTAAATCA
CATCATTACAACTCATCATCCTGTCGCCATAGCGGAGGAAGCTTGCTTATTGGATCAGT
TAAGTGAAGGGAGATTTATTTTAGGGTTTAGTGATTGCGAAAAAAAAGATGAAATGCATT
TTTTTAATCGCCCGGTTGAATATCAACAGCAACTATTTGAAGAGTGTTATGAAATCATTA
ACGATGCTTTAACAACAGGCTATTGTAATCCAGATAACGATTTTTATAGCTTCCCTAAAA
TATCTGTAAATCCCCATGCTTATACGCCAGGCGGACCTCGGAAATATGTAACAGCAACC
AGTCATCATATTGTTGAGTGGGCGGCCAAAAAAGGTATTCCTCTCATCTTTAAGTGGGA
TGATTCTAATGATGTTAGATATGAATATGCTGAAAGATATAAAGCCGTTGCGGATAAATA
TGACGTTGACCTATCAGAGATAGACCATCAGTTAATGATATTAGTTAACTATAACGAAGA
TAGTAATAAAGCTAAACAAGAGACGCGTGCATTTATTAGTGATTATGTTCTTGAAATGCA
CCCTAATGAAAATTTCGAAAATAAACTTGAAGAAATAATTGCAGAAAACGCTGTCGGAAA
TTATACGGAGTGTATAACTGCGGCTAAGTTGGCAATTGAAAAGTGTGGTGCGAAAAGTG

TATTGCTGTCCTTTGAACCAATGAATGATTTGATGAGCCAAAAAAATGTAATCAATATTG
TTGATGATAATATTAAGAAGTACCACATGGAATATACCTAATAGATTTCGAGTTGCAGCG
AGGCGGCAAGTGAACGAATCCCCAGGAGCATAGATAACTATGTGACTGGGGTGAGTGA
AAGCAGCCAACAAAGCAGCAGCTTGAAAGATGAAGGGTATAAAAGAGTATGACAGCAG
TGCTGCCATACTTTCTAATATTATCTTGAGGAGTAAAACAGGTATGACTTCATATGTTGA
TAAACAAGAAATTACAGCAAGCTCAGAAATTGATGATTTGATTTTTTCGAGCGATCCATT
AGTGTGGTCTTACGACGAGCAGGAAAAAATCAGAAAGAAACTTGTGCTTGATGCATTTC
GTAATCATTATAAACATTGTCGAGAATATCGTCACTACTGTCAGGCACACAAAGTAGATG
ACAATATTACGGAAATTGATGACATACCTGTATTCCCAACATCGGTTTTTAAGTTTACTC
GCTTATTAACTTCTCAGGAAAACGAGATTGAAAGTTGGTTTACCAGTAGCGGCACGAAT
GGTTTAAAAAGTCAGGTGGCGCGTGACAGATTAAGTATTGAGAGACTCTTAGGCTCTGT
GAGTTATGGCATGAAATATGTTGGTAGTTGGTTTGATCATCAAATAGAATTAGTCAATTT
GGGACCAGATAGATTTAATGCTCATAATATTTGGTTTAAATATGTTATGAGTTTGGTGGA
ATTGTTATATCCTACGACATTTACCGTAACAGAAGAACGAATAGATTTTGTTAAAACATT
GAATAGTCTTGAACGAATAAAAAATCAAGGGAAAGATCTTTGTCTTATTGGTTCGCCATA
CTTTATTTATTTACTCTGCCATTATATGAAAGATAAAAAAATCTCATTTTCTGGAGATAAA
AGCCTTTATATCATAACCGGAGGCGGCTGGAAAAGTTACGAAAAAGAATCTCTGAAACG
TGATGATTTCAATCATCTTTTATTTGATACTTTCAATCTCAGTGATATTAGTCAGATCCGA
GATATATTTAATCAAGTTGAACTCAACACTTGTTTCTTTGAGGATGAAATGCAGCGTAAA
CATGTTCCGCCGTGGGTATATGCGCGAGCGCTTGATCCTGAAACGTTGAAACCTGTAC
CTGATGGAACGCCGGGGTTGATGAGTTATATGGATGCGTCAGCAACCAGTTATCCAGC
ATTTATTGTTACCGATGATGTCGGGATAATTAGCAGAGAATATGGTAAGTATCCCGGCG
TGCTCGTTGAAATTTTACGTCGCGTCAATACGAGGACGCAGAAAGGGTGTGCTTTAAGC
TTAACCGAAGCGTTTGATAGTTGA

**RBS30**ATTAAAGAGGAGAAA

**RBS35**ATTAAAGAGGAGAA

**RBS64**AAAGAGGGGAAA

**Bxb1B** CGGCCGGCTTGTCGACGACGGCGGTCTCCGTCGTCAGGATCATCCGGGC

**Bxb1P**
GTCGTGGTTTGTCTGGTCAACCACCGCGGTCTCAGTGGTGTACGGTACAAACC
CCGAC

**PnorV**
ACGGAAAAACTCATCTTTGCCTCACTGTCAATTTGACTATAGATATTGTCATATC
GACCATTTGATTGATAGTCATTTTGACTACTCATTAATGGGCATAATTTTATTTATAGAGT
AAAAACAATCAGATAAAAAACTGGCACGCAATCTGCAATTAGCAAGACATCTTTTTAGAA
CACG

**NorR**
ATGAGTTTTTCCGTTGATGTGCTGGCGAATATCGCCATCGAATTGCAGCGTGGG
ATTGGTCACCAGGATCGTTTTCAGCGCCTGATCACCACGCTACGTCAGGTGCTGGAGT
GCGATGCGTCTGCGTTGCTACGTTACGATTCGCGGCAGTTTATTCCGCTTGCCATCGA
CGGTCTGGCAAAGGATGTACTCGGTAGACGCTTTGCGCTGGAAGGGCATCCACGGCT
GGAAGCGATTGCCCGCGCCGGGGATGTGGTGCGCTTTCCCGCAGACAGCGAATTGCC
CGATCCCTATGACGGTTTGATTCCTGGGCAGGAGAGTCTGAAGGTTCACGCCTGCGTT
GGTCTGCCATTGTTTGCCGGGCAAAACCTGATCGGCGCACTGACGCTCGACGGGATG
CAGCCCGATCAGTTCGATGTTTTCAGCGACGAAGAGCTACGGCTGATTGCTGCGCTGG
CGGCGGGAGCGTTAAGCAATGCGTTGCTGATTGAACAACTGGAAAGCCAGAATATGCT
GCCAGGCGATGCCACGCCGTTTGAAGCGGTGAAACAGACGCAGATGATTGGCTTGTCC

CCTGGCATGACGCAACTGAAAAAAGAGATTGAGATTGTGGCGGCGTCCGATCTCAACG
TCCTGATCAGCGGTGAGACTGGAACCGGTAAGGAGCTGGTGGCGAAAGCGATTCATGA
AGCCTCGCCACGGGCGGTGAATCCGCTGGTCTATCTCAACTGTGCTGCACTGCTGGAA
AGTGTGGCGGAAAGTGAGTTGTTCGGGCATGTGAAAGGAGCGTATACTGGCGCTATCA
GTAATCGCAGCGGGAAGTTTGAAATGGCGGATAACGGCACGCTGTTTCTGGATGAGAT
CGGCGAGTTGTCGTTGGCATCGCAGGCCAAGCTGCTGAGGGTGTTGCAGTATGGCGA
TATTCAGCGCGTTGGCGATGACCGTTGTTTGCGGGTCGATGTGCGCGTGCTGGCGGC
GACTAACCGCGATTTACGCGAAGAGGTGCTGGCAGGGCGATTCCGCGCCGATTTGTTT
CATCGCCTGAGCGTGTTTCCACTTTCGGTGCCGCCGCTGCGTGAGCGGGGCGATGAT
GTCATTCTGCTGGCGGGGTATTTCTGCGAGCAGTGTCGTTTGCGGCAGGGGCTCTCCC
GCGTGGTATTAAGTGCCGGAGCGCGAAATTTACTGCAACACTACAGTTTTCCGGGAAA
CGTGCGCGAACTGGAACATGCTATTCATCGGGCGGTAGTTCTGGCGAGAGCCACCCG
CAGCGGCGATGAAGTGATTCTTGAGGCGCAACATTTTGCGCGCGCGGATGCTGGAAAC
CGACGTCGCCAACCTGCATCGGCTGGCGAAACGCGCGCGGATGCTGGAAACGCGACA
GAAGCGTTCCAGCGTGAAACTATTCGTCAGGCACTGGCACAAAATCATCACAACTGGG
CTGCCTGCGCGCGGATGCTGGAAACCGACGTCGCCAACCTGCATCGGCTGGCGAAAC
GCGCGCGGATGCTGGAAACCGACGTCGCCAACCTGCATCGGCTGGCGAAACGTCTGG
GATTGAAGGATTAA

**TtrR**

ATGAGCCTGGCGCTGCCGGTGTACCTGATTGACGACGATGACTCCGTTCGCCG
TTCCCTGCGTTTCATGCTGGAAAGCTACGGCCTGAAAATCACTGACTTCGATTCTGCTG
AAGCGTTCTTCACCGCGGTAGACCTGACCCTGCCGGGTTGCGCACTGGTGGACGTAC
GTATGCCGGGCCTGAGCGGCCCACAGCTGCACCTGGAACTGGTGTCTAAGAACAGCC
CGCTGGCCGTGATCTATCTGACCGGTCACGGCGACGTTCCGATGGCGGTTGAAGCGC
TGAAACTGGGTGCGGTAGATTTCTTTCAGAAACCTGCAGACGGCGCTAAACTGGCTGA
AGCTGTGGTCAAAGCCCTGGAACACGCGAAAACCCACTACCAAGACAACCAGTACCTG
GAAACCTATCAGGCTCTGACCCCACGTGAGCGCGAAATCCTGAACCTGATTGCGCAAG
GTCTGAAAAACCAGGAAATCGCGGACAACCTGTGCATTGCGATGCGCACCGTAGAAAT
CCACCGTGCGAACCTGATGAAAGGTATGCAGGTGGGCAGCCTGGCAGAGATGATGCT
GATTTACTCTCGTATCGCGGAGCGTCTGCGCCTGGAAGATAACCGTTAA

**TtrS**

ATGTTTTCTTCCGGCATGAGCCTGAAATGGCCTCTGAGCATCCAGGGTTCCATC
GAACCGAAAATGACTCCGCGCATCATCTTCCGCATGGCCCTGCGCACTAAAGTGTTCAT
CACCCTGCTGGCTTGCCTGTTTGCTTCTGCCAACGTATTCGCGGTGGAACCGCAGAAC
CAGTCCGCGCCTTCCGATGTGATGGACGCTGAGGTGGTTGCTCCGGTTGCGACGGAG
GAACCGGGTTACCAGGTCGTTGATGTTGGTGTGCTGGCGATTCGCGGTTATAAAGCTA
CCATCAACCGTTGGCAGCCACTGATGGGTTGGCTGGAGACTCAGATCCCGAACTCCTA
CTTCCGCCTGCACCCGCTGTCTCTGGACGAGCTGGCAAAGGGTGTAGAAACCCAGGG
TCTGGACTTCGTAATCACCAATCCAGGCCAGTCCGTACTGCTGGCGCGTCAGTATTCTC
TGACGTGGCTGGCGACCCTGCGCTCTCCGCTGAACAACGGTGCTGCGATGCAGGTGG
GTTCCGCGCTGGTAGTTCGTGCCGACAGCCCGTACCAGACCCTGACCGATCTGAAAGG
CCGTCGTCTGGGCATCGTCAGCAAAAACGCATTCGGTGGTTATCTGACGCTGGTATAT
GAGGCACAGCTGAAAGGCATCGATCTGCCGCGCTTTGTTGGTGAAATCATCCCACTGG
GCTTCCCGCTGGATAACCTGCTGTATCAGCTGGATGATCAGAAATCTGGTAATGACGCC
GCAAAAGACAATCGTCTGGATGCCACCGTTGTACCGGTATGCCAACTGGAACAGATGC
AGGCCGAAGGTCTGATCAACATCGGCCACTATCGCGTTCTGGACAACCAGACCCCGGT
AGGTTTCCACTGTCAAGTAAGCACTCGTCTGTATCCGAACTGGTCTATGGCTAAAACTA
ACCGTGCCAGCCAGTCCCTGGCAAAATCTGTGACCCAGGCTCTGCTGGCTCTGCCGGA
AGATCACCTGGCAGCGAAAAGCTCTGACTCTGCAGGTTGGACCACTGCAGTGTCCCAG
CTGGCGATCGACCAGCTGCTGAAAGACCTGGACATGCATCCACTGCAGACCCCATGGT

GGCAGCGCGCTTGGCAGTGGGTAAAACTGCACCAGCAGTGGGCGTGGTTCATTCTGG
GTATCCTGGTCCTGCTGAACGCATACCATTTCTGGCTGGAATACCGTTTCTCCCGTCGT
GGTCGTGAACTGATCAACACCCAGCGCCAGCTGAACGAAAACCGTGCTCTGCTGGAGC
ACGCACAACGTATCGCCATCGCTGGCGAACTGGGTGCTAGCCTGTCCCATGAACTGAA
CCAGCCGCTGGCTGCAATTGGTCACTATTGCCATGGTGCGGAAGTGCGTCTGCAGCGT
GGTACTTCTCCTGAAGAACTGCAGTCTGTTCTGACCCTGATCCAGCAGGAAGTTACGC
GCGCCGACTCTATCATCAGCCGTCTGCGTAACCTGCTGAAAAAACGCCCGGTTAGCAA
ACAGGCTCTGTATCTGCACGAACTGGTAAACGAAACGGTGCCGCTGCTGGCGTACGAA
TTCGAACAGCACCAGATCAACCTGGCGGTAAACGTCAGCGGTGAACCTTACCTGCAAT
CTCTGGACGAGGTCGGTATGCAGCAGCTGCTGCTGAACCTGCTGAAAAACGCGTTTGA
CGCGTGCGTTCAGCGCCTGGAACTGGAATCCTCCGGCACTGAACAGAGCGGTATGCG
TAAACCGTATGTTCCTACGATCGATATTGATCTGCGCTACCAGGAGCGTACTCTGCTGC
TGACTGTGACGGATAACGGCACTGGTCTGACCGAGGAGACTAGCCTGCTGATGCAGG
CCTTCTATTCCACCAAATCTGAAGGCCTGGGCCTGGGTCTGGTTATTTGCCGTGATATC
GCTGAGTCCCACGGTGGCACCTTCTCTCTGGAATCCGCCATGGGTGGTGGCTGCCAG
GCTCAGGTGGCCATCCCGCGCAAGCCGGAACCGAATGGCGTTCTGTAA

**TtrB**

TTTATAGTAAATCACTGCATAATTCGTGTCGCTCAAGGCGCACTCCCGTTCTGGA
TAATGTTTTTTGCGCCGACATCATAACGGTTCTGGCAAATATTCTGAAATGAGCTGGTTA
GCTAGTCAATATTTGTTGCGCTAGATCAAATCCACGCCTGATATGTGGAAAACCACTATA
GTTATGCCGCCTCGCCTTTTACAGAATGCCTGT
